# CD40 ligand and IFN-γ induce proinflammatory activity and microglia-mediated neurodegeneration in human stem cell models

**DOI:** 10.64898/2026.10.28.749776

**Authors:** Gregory S. Wirak, Sean K. Ryan, Anna Blazier, Dongyu Liu, Ivan Nyarko-Danquah, Michael LaMorte, Shameer Khader, Jose Vela, Bo Yang, Hendrik Wesseling, Bailin Zhang, Franck Rapaport, Roopali Gandhi, Biljana Djukic, Dimitry Ofengeim, Timothy R. Hammond

**Author notes:** Employee of Sanofi when this work was undertaken (Gregory S. Wirak, Roopali Gandhi, Dimitry Ofengeim). These authors contributed equally to this work (Gregory S. Wirak, Sean K. Ryan). **Corresponding author:** Timothy Hammond, Address: 350 Water Street, Cambridge, MA 02141.

## Abstract

The cluster of differentiation 40 (CD40)/CD40 ligand (CD40L) co-stimulatory pathway plays a critical role in the activation and regulation of immune networks involving both adaptive and innate immune cells. Dysregulation of this pathway has been implicated in multiple autoimmune diseases, including multiple sclerosis (MS). MS is characterized by progressive neurodegeneration driven by compartmentalized neuroinflammation within the central nervous system (CNS), sustained low-grade T cell infiltration at the blood-brain barrier, and T cell reactivation at CNS border tissues. CD40L-expressing T cells can activate CD40-expressing microglia, which are a critical effector cell that contribute to neurodegeneration in MS. However, the consequence of this functional interaction in the brain remains poorly understood. Here, we demonstrate that the combination of CD40L and interferon-gamma (IFN-γ), a cytokine produced by activated T cells, induces a robust inflammatory response and cell death in human induced pluripotent stem cells (hiPSC)-derived microglia. Stimulation with CD40L elicited microglial clustering, morphological transformation, and robust release of proinflammatory cytokines including interleukin-12 (IL-12), tumor necrosis factor-alpha (TNF-α), and IL-6. Anti-CD40L antibody treatment reversed both cell death and inflammatory response and partially normalized the associated transcriptional changes. Furthermore, using an hiPSC-derived tri-culture system comprising neurons, astrocytes, and microglia, we demonstrated that CD40L and IFN-γ trigger microglia-dependent neuronal dysregulation and death. Anti-CD40L treatment substantially attenuated neuronal injury. Finally, transcriptomic enrichment analysis identified CD40L-mediated microglial gene expression signatures that correlated with CD40 expression in primary progressive MS brain tissue but not in the normal-appearing white matter of healthy controls. Collectively, these findings support a role for the microglial CD40/T cell CD40L axis in MS pathology and suggest CD40L blockade as a potential therapeutic strategy warranting further investigation.

## INTRODUCTION

Multiple sclerosis (MS) is a chronic, immune-mediated, neurodegenerative disease of the central nervous system (CNS) characterized by demyelination and progressive neurodegeneration^1,2^. Dysregulation of co-stimulatory pathways has been implicated in multiple autoimmune diseases^3,4^, including MS, in which genetic variants near CD40 are associated with MS risk^5^, and immune cell infiltration of the CNS, including T cells, promotes inflammation, demyelination and axon degeneration^6^. Despite the availability of >20 disease-modifying therapies, most of which primarily target relapses via the peripheral immune system for relapsing-remitting MS (RRMS), most patients inevitably experience disease progression over time and transition to secondary progressive MS (SPMS)^7,8^. Emerging evidence indicates that relapse suppression does not correlate well with attenuation of disease progression^9,10^, highlighting a critical unmet need for therapies that target both peripheral adaptive immunity and CNS-resident innate immune cells.

Aberrant cluster of differentiation 40 (CD40)/CD40 ligand (CD40L) signaling drives pathogenic immune responses, contributing to chronic inflammation and tissue damage^11^. In MS, the CD40/CD40L co-stimulatory axis represents a mechanistic bridge linking peripheral adaptive immunity and CNS innate immune activation^12^. CD40, a member of the tumor necrosis factor (TNF) receptor superfamily, is expressed on antigen-presenting cells including B cells, dendritic cells, monocytes, and microglia^3,4^. Its cognate ligand, CD40L (also known as CD154), is predominantly expressed on activated CD4^+^ T cells but also found on activated platelets, endothelial cells, and other immune cells^13^. In the periphery, CD40L-expressing T cells engage CD40 on B cells, triggering cellular activation, antibody production, and proinflammatory cytokine secretion^4^. In MS, these activated T cells and B cells can then traverse the blood-brain barrier (BBB) and enter the brain parenchyma, where they contribute to lesion formation and neuroaxonal damage^2,6^. Microglial CD40 activation has been implicated in MS where microglia express CD40 and can be activated by CD40L-expressing T cells^12,14^. CD40 deficiency in CNS-resident microglia attenuated full microglial activation and ameliorated experimental autoimmune encephalomyelitis (EAE) severity in bone marrow chimeric mice^15^. CD40L^+^ cells are also found in the parenchyma of patients with MS and co-localize with CD40^+^ cells in active lesions^12^. Interferon-gamma (IFN-γ), a proinflammatory cytokine produced by activated T cells and elevated in the cerebrospinal fluid (CSF) of patients with MS^16,17^, significantly upregulates microglial CD40 expression and amplifies CD40L-dependent activation^14,18,19^. Despite this convergent evidence implicating the microglial CD40/T cell CD40L axis in MS pathology, the direct impact of this interaction on microglial inflammatory responses and neuronal viability remains incompletely defined.

In this study, we investigated the effects of stimulation with CD40L on human induced pluripotent stem cell (hiPSC)-derived microglia and assessed the therapeutic potential of CD40L blockade. We demonstrate that CD40L elicits a robust inflammatory response and cell death in hiPSC-derived microglia, an effect that is further exacerbated by co-stimulation with IFN-γ. Importantly, anti-CD40L treatment effectively abolishes this response. Bulk RNA sequencing of stimulated microglia revealed activation of multiple immune-related pathways, including modules linked to B cell activation and proliferation. Employing a hiPSC-derived tri-culture system (microglia, neurons, and astrocytes), we further demonstrate that co-stimulation with CD40L and IFN-γ induces significant reduction in neuronal activity, which is partially rescued by anti-CD40L treatment. Finally, we demonstrate that the CD40L-induced microglial gene signature selectively correlates with CD40 expression in the microglia isolated from perilesional brain tissue of patients with primary progressive MS (PPMS) but not from normal-appearing white matter (NAWM) of healthy controls. Collectively, these findings provide evidence that microglial CD40/T cell-CD40L interactions drive CNS-intrinsic inflammation and highlight CD40L blockade as a potential therapeutic avenue in MS.

## RESULTS

### Co-stimulation of human microglia with CD40L and IFN-γ elicits cell clustering, ramification, cytokine release, and cell death

We first examined the morphological and behavioral responsiveness of hiPSC-derived microglia to CD40L. Stimulation with CD40L alone, or in combination with IFN-γ, elicited pronounced morphological changes 24 h after stimulation. Microglia stimulated with CD40L alone formed clusters and adopted a unidimensional elongated morphology, whereas co-stimulation with CD40L and IFN-γ resulted in more extensive clustering and a more ramified morphology (Fig. 1A and 1B and Fig. S1 and S2). Treatment with a CD40L-targeting antibody reduced clustering under both conditions (Fig. S1). Quantification of the nearest-neighbor nuclear distances confirmed that stimulation with CD40L elicited microglial clustering, an effect that was blocked by anti-CD40L antibody treatment. Co-stimulation with CD40L and IFN-γ induced an even greater degree of clustering, which was likewise partially rescued by anti-CD40L treatment (Fig. 1C and Fig. S2).

**Fig. 1.**
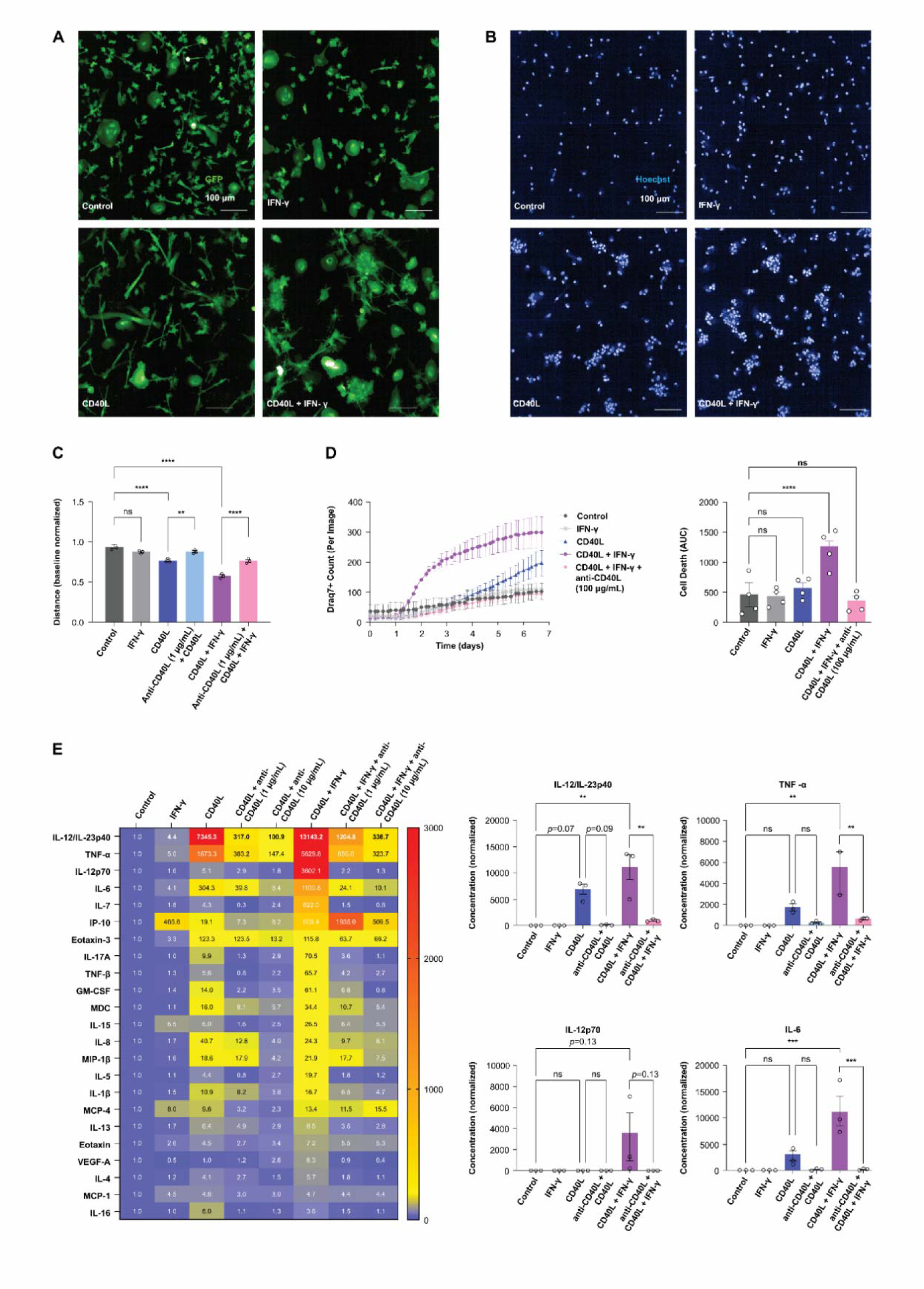
Co-stimulation of CD40L and IFN-γ on hiPSC-derived microglia drives inflammation and cell death. A Representative confocal images of live GFP-expressing hiPSC-derived microglia at 24 h post co-stimulation with CD40L and/or IFN-γ. B Representative confocal images of Hoechst-stained nuclei in live microglia at 24 h post co-stimulation with CD40L and/or IFN-γ. C Nuclei nearest-neighbor distance quantified from confocal fluorescence images of Hoechst-stained microglia (as in B) at 24 h post stimulation with CD40L, IFN-γ, and/or an anti-CD40L tool antibody (n=3 independent experiments). D Time course of dead-cell counts in DRAQ7-stained microglia cultures up to 7 days post stimulation with CD40L, IFN-γ, and/or an anti-CD40L tool antibody (n=3 independent experiments). E Cytokine concentrations in culture supernatants from microglia co-stimulated with CD40L, IFN-γ, and/or an anti-CD40L tool antibody. The heatmap (left panel) denotes average cytokine levels across experiments, normalized to vehicle control within each experiment. Bar charts (right panel) show average cytokine levels across experiments, normalized to vehicle control within experiment, with each dot representing an individual experiment (n=3 independent experiments). Statistical significance is indicated as * *p*-value≤0.05, ** *p*-value≤0.01, *** *p*-value≤0.001, and **** *p*-value≤0.0001. Data are presented as mean±SEM. CD40L, cluster of differentiation 40 ligand; DRAQ7, deep red anthraquinone 7; GFP, green fluorescent protein; hiPSC, human induced pluripotent stem cell; IFN-γ, interferon-gamma; SEM, standard error of the mean.

To assess viability following co-stimulation with CD40L and/or IFN-γ, we quantified deep red anthraquinone 7 (DRAQ7)-positive dead cells over 7 days post stimulation. Control and IFN-γ-stimulated cultures exhibited minimal cell death over the examined period. In contrast, cultures exposed to co-stimulation of CD40L and IFN-γ triggered a progressive increase in cell death, beginning approximately 36 h post stimulation, with a faster rate of cell death than with CD40L alone. CD40L alone induced cell death only at later timepoints without achieving significance, while anti-CD40L treatment reduced cell death under co-stimulation conditions (Fig. 1D). Inhibitors of necroptosis (RIPK1i, RIPK3i), pyroptosis (NLR family pyrin domain-containing 3 (NLRP3i)), and ferroptosis (liproxstatin-1) did not prevent CD40L+IFN-γ-induced cell death (Fig. S3), suggesting that an alternative form of cell death occurs in these cells. To characterize microglial inflammatory responses to co-stimulation with CD40L and/or IFN-γ, we performed multiplex cytokine profiling on cell culture supernatants collected 24 h post stimulation to avoid the confounding effects of cell death. Across 23 analytes, CD40L elicited a robust cytokine release profile, which was broadly enhanced by co-stimulation with IFN-γ. IFN-γ alone triggered a comparatively modest response, although it did increase the release of multiple cytokines, most notably IFN-γ-induced protein 10 (IP-10) (Fig. 1E, heatmap). The four cytokines most strongly induced by co-stimulation with CD40L and IFN-γ were interleukin-12 (IL-12)/IL-23p40, tumor necrosis factor-alpha (TNF-α), IL-12p70, and IL-6 (Fig. 1E, bar graphs and Fig. S4). For these and other cytokines, co-stimulation produced levels that exceeded the simple additive effects of CD40L alone and IFN-γ alone.

### Co-stimulation of human microglia with CD40L and IFN-γ elicits an inflammatory transcriptional response

To gain greater insight into the transcriptional consequences of CD40L signaling in human microglia, we co-stimulated hiPSC-derived microglia with CD40L and/or IFN-γ and performed bulk RNA sequencing. Principal component analysis (PCA) revealed a clear separation between treatment conditions with anti-CD40L partially reversing transcriptomic changes in both CD40L and CD40L + IFN-γ conditions (Fig. 2A).

**Fig. 2.**
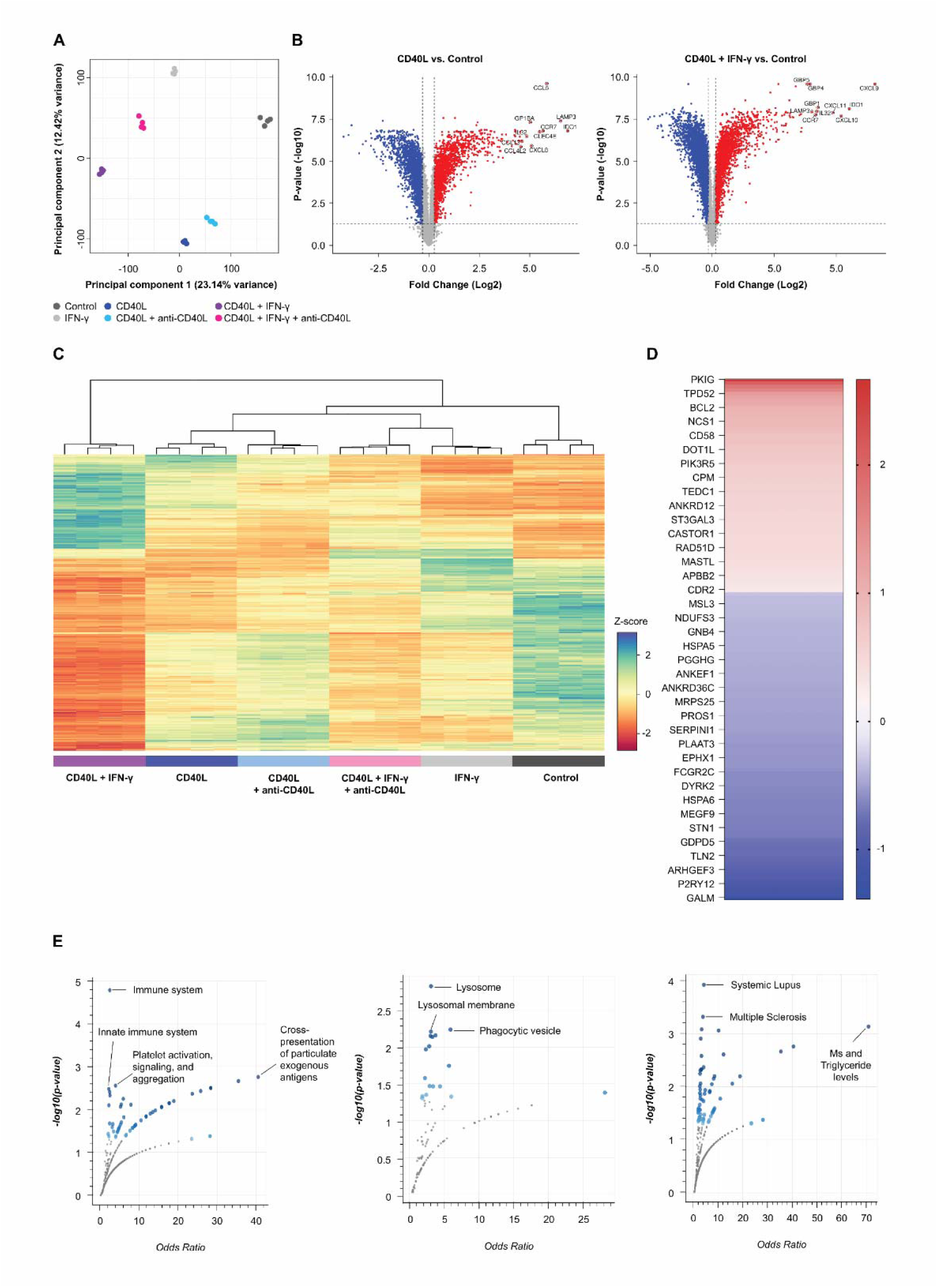
CD40L and IFN-γ co-stimulated microglia acquire an inflammatory, disease-associated transcriptomic profile. A PCA plot showing the first two principal components calculated from bulk RNA sequencing of hiPSC-derived microglial cultures, 12 h post co-stimulation with CD40L, IFN-γ, and/or an anti-CD40L tool antibody (n=4 technical replicates). B Volcano plots of differential gene expression comparing CD40L-stimulated vs. unstimulated microglia (left panel) and CD40L and IFN-γ co-stimulated vs. unstimulated microglia (right panel), within bulk RNA sequencing dataset described in A, (|log2 fold change|>1 (≥2-fold), FDR≤0.05)). C Hierarchical clustering of conditions by gene expression. Heatmap displays DESeq2 normalized counts scaled using a Z-score. D Heatmap of genes that were upregulated or downregulated in the CD40L+IFN-γ condition that were significantly reversed by anti-CD40L treatment plotted by fold change compared to control. E Pathway analysis of reversed genes. Panels from left to right, top to bottom: Reactome, Ontologies cellular component, Disease/drugs GWAS catalog. All pathway analysis performed through enrichr.com. CD40L, cluster of differentiation 40 ligand; DESeq2, Differential Expression analysis based on the negative binomial distribution (version 2); FDR, false discovery rate; GWAS, Genome-Wide Association Study; hiPSC, human induced pluripotent stem cell; *IDO1*, indoleamine 2,3-dioxygenase 1; IFN-γ, interferon-gamma; PCA, principal component analysis; RNA, ribonucleic acid.

Differential expression analysis revealed that, compared to the control, stimulation with CD40L induced upregulation of multiple inflammatory and antigen-presentation-related genes. Among the most strongly upregulated transcripts in CD40L-stimulated microglia were *CCL5, GP1BA, LAMP3, IDO1, CCR7, CLEC4E,* and *CXCL8*, while a broad set of genes also showed reduced expression (Fig. 2B, left panel). Co-stimulation with CD40L and IFN-γ further amplified this response and induced additional interferon-responsive genes, with the most strongly upregulated transcripts being *GBP1, GBP4, GBP5, CXCL9, CXCL10, CXCL11, IL32, LAMP3, IDO1,* and *CCR7* (Fig. 2B, right panel). Consistent with our cytokine profiling, many transcripts exhibited a greater magnitude of change with co-stimulation than with either stimulus alone. Anti-CD40L antibody treatment broadly attenuated these transcriptional responses, reducing the expression of CD40L-responsive and IFN-γ-enhanced genes (Fig. 2C).

To determine which genes were specifically regulated by CD40L, we identified a subset of dysregulated genes in the combined CD40L and IFN-γ condition that were fully reversed to control levels by anti-CD40L treatment (Fig. 2D). Pathway analysis of these specific genes identified multiple inflammatory, lysosomal, and phagocytic-related pathways that were controlled by CD40L. (Fig. 2E). Dysregulation of lysosomal lipid processing and phagocytic clearance are established features in MS lesion-associated microglia and macrophages^20,21^, suggesting that CD40L-mediated activation may further compromise these critical homeostatic processes. These findings suggest that CD40L-induced microglia activation produces disease-associated transcriptomic changes that may contribute to progression.

### Human microglia co-stimulated with CD40L and IFN-γ mediate neuronal injury

Smoldering neuroinflammation can drive neuronal damage and disease progression in MS^9^. Potential cellular sources of this inflammation include resident microglia, invading macrophages, B cells, and/or T cells^22^. We hypothesized that T cells present in CNS lesions, which express CD40L^12^, may chronically activate microglia and contribute to neuronal damage; to test this hypothesis, we utilized an all-hiPSC-derived tri-culture comprising neurons, astrocytes, and microglia^23^. Neuronal function within the tri-cultures was assessed via live calcium imaging of active neurons. Co-stimulation with CD40L and IFN-γ resulted in a pronounced reduction in active neuron counts over the 11-day observation period (Fig. 3A and 3B).

**Fig. 3.**
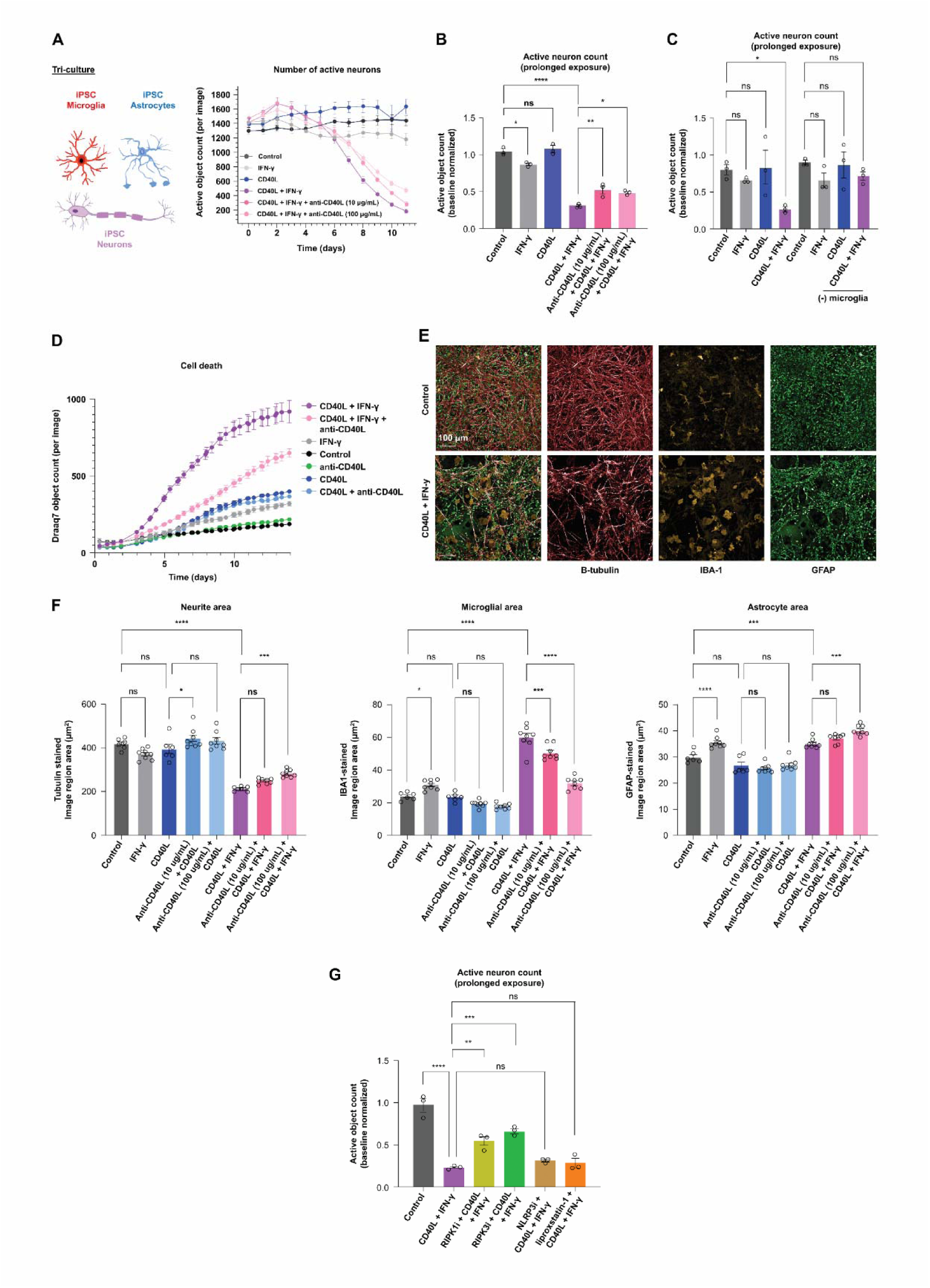
Microglia-mediated neuronal insult in a hiPSC-derived tri-culture co-stimulated with CD40L and IFN-γ. A Time course of active neuron counts within hiPSC-derived tri-cultures of microglia, astrocytes, and neurons, up to 11 days after co-stimulation with CD40L, IFN-γ, and/or an anti-CD40L tool antibody (n=6 technical replicates). Displayed are average counts per image, with one time-lapse image acquired per culture-plate well. Neuronal activity was captured via neuron-specific calcium imaging every 24 h. Active neuron counts within tri-cultures or co-cultures of astrocytes and neurons co-stimulated with B an anti-CD40L tool antibody and/or with C CD40L, IFN-γ, (n=3 independent experiments per graph, as in A). Counts were quantified across conditions at the first timepoint (assessed every 24 h) at which the average count under the co-stimulated with CD40L and IFN-γ condition decreased by ≥70% relative to baseline (i.e., time 0 in A). Values shown are active neuron counts normalized to within-condition baseline counts. D Time course of dead cell counts in DRAQ7-stained tri-cultures, up to 14 days after co-stimulation with CD40L, IFN-γ, and/or an anti-CD40L tool antibody (n=6 technical replicates). E Fluorescent immunostaining for markers of neurons (beta-III tubulin), microglia (IBA1), and astrocytes (GFAP) in tri-cultures co-stimulated with CD40L and IFN-γ at 14 days after stimulation. F Neurite, microglial, and astrocyte areas in tri-cultures co-stimulated with CD40L, IFN-γ, and/or an anti-CD40L tool antibody, as assessed by the immunostaining in E. G Active neuronal object counts 11-days post-treatment with recombinant human IFN-γ (100 U/mL), recombinant human CD40L (5 μg/mL), RIPK1 inhibitor GSK547 (10 μM), RIPK3 inhibitor GSK872 (3 μM), ferroptosis inhibitor liproxstatin-1 (100 nM), and/or the NLRP3 inflammasome inhibitor MCC950 (5 μM). Bars denote the average stained image region area in µm^2^ (n=6–8 technical replicates). Statistical significance is indicated as * *p*-value≤0.05, ** *p*-value≤0.01, *** *p*-value≤0.001, and **** *p*-value≤0.0001. Data are presented as mean±SEM. CD40L, cluster of differentiation 40 ligand; DRAQ7, deep red anthraquinone 7; GFAP, glial fibrillary acidic protein; hiPSC, human induced pluripotent stem cell; IBA1, ionized calcium-binding adapter molecule 1; IFN-γ, interferon-gamma; SEM, standard error of the mean.

Co-treatment with a CD40L-targeting antibody reduced, but did not fully restore, the loss of active neurons associated with co-stimulation with CD40L and IFN-γ (Fig. 3A and 3B). Importantly, cultures lacking microglia did not exhibit changes in neuronal activity under any condition, suggesting that microglia are necessary to induce the neuronal dysfunction associated with co-stimulation with CD40L and IFN-γ (Fig. 3C). Over a 14-day period, co-stimulation with CD40L and IFN-γ also caused marked cell death in DRAQ7-stained tri-cultures, which was reduced by CD40L antibody co-treatment (Fig. 3D).

Immunocytochemistry at day 14 revealed that, relative to control cultures, co-stimulation with CD40L and IFN-γ significantly reduced neurite density (Fig. 3E and F). Neither CD40L nor IFN-γ alone elicited this effect, and co-treatment with a CD40L-targeting antibody partially attenuated the loss of neurite complexity (Fig. 3F). Under co-stimulation conditions, microglia adopted an enlarged amoeboid morphology accompanied by a pronounced increase in microglial ionized calcium-binding adapter molecule 1 (IBA1) area. IFN-γ alone minimally increased microglial IBA1-positive area, but this effect was far smaller than that elicited by CD40L and IFN-γ in combination (Fig. 3E and F). These morphological and staining area changes were reversed by CD40L blockade (Fig. 3F, Fig. S6, and Fig. S7). Unlike CD40L, IFN-γ alone increased astrocytic glial fibrillary acidic protein (GFAP) immunostaining area, but the effect of co-stimulation was comparable to that of IFN-γ alone, suggesting that CD40L activation specifically affects microglia (Fig. 3F). While neither the ferroptosis inhibitor liproxstatin-1 nor an NLRP3 inflammasome inhibitor modulated active neuron loss, inhibition of RIPK1 or RIPK3 significantly reduced the loss of active neurons (Fig. 3G), suggesting the involvement of necroptosis.

To investigate how microglial morphology evolves over time in this tri-culture system following co-stimulation with CD40L and/or IFN-γ, we utilized GFP-expressing microglia and performed longitudinal live-cell imaging over 10 days. In parallel, active neuronal populations were quantified via calcium imaging, enabling correlation of microglial morphological dynamics with the progressive loss of functional neurons (Fig. S8A). As observed in microglia mono-cultures (Fig. 1C), both stimulation with CD40L alone and co-stimulation with CD40L and IFN-γ elicited microglial clustering after 24 h. Notably, co-stimulation additionally drove the emergence of a hyper-ramified morphology at this same timepoint, consistent with mono-culture findings (Fig. 1B). Over the subsequent 7–10 days, both clustering and hyper-ramification gradually diminished, and microglia exposed to CD40L and IFN-γ adopted an enlarged amoeboid phenotype, coinciding with the greatest reductions in active neuron counts (Fig. S8B).

We next profiled the cytokines released into the tri-culture medium upon co-stimulation with CD40L and/or IFN-γ. Cytokine profiling at 24 h post stimulation and at 5 days post stimulation (Fig. 4A and 4B) showed that at both timepoints, stimulation with CD40L alone elicited a robust cytokine release, which was broadly enhanced by co-stimulation with IFN-γ. Among the most strongly elevated cytokines under co-stimulation conditions were IL-12/IL-23p40, TNF-α, IL-12p70, and IL-6.

**Fig. 4.**
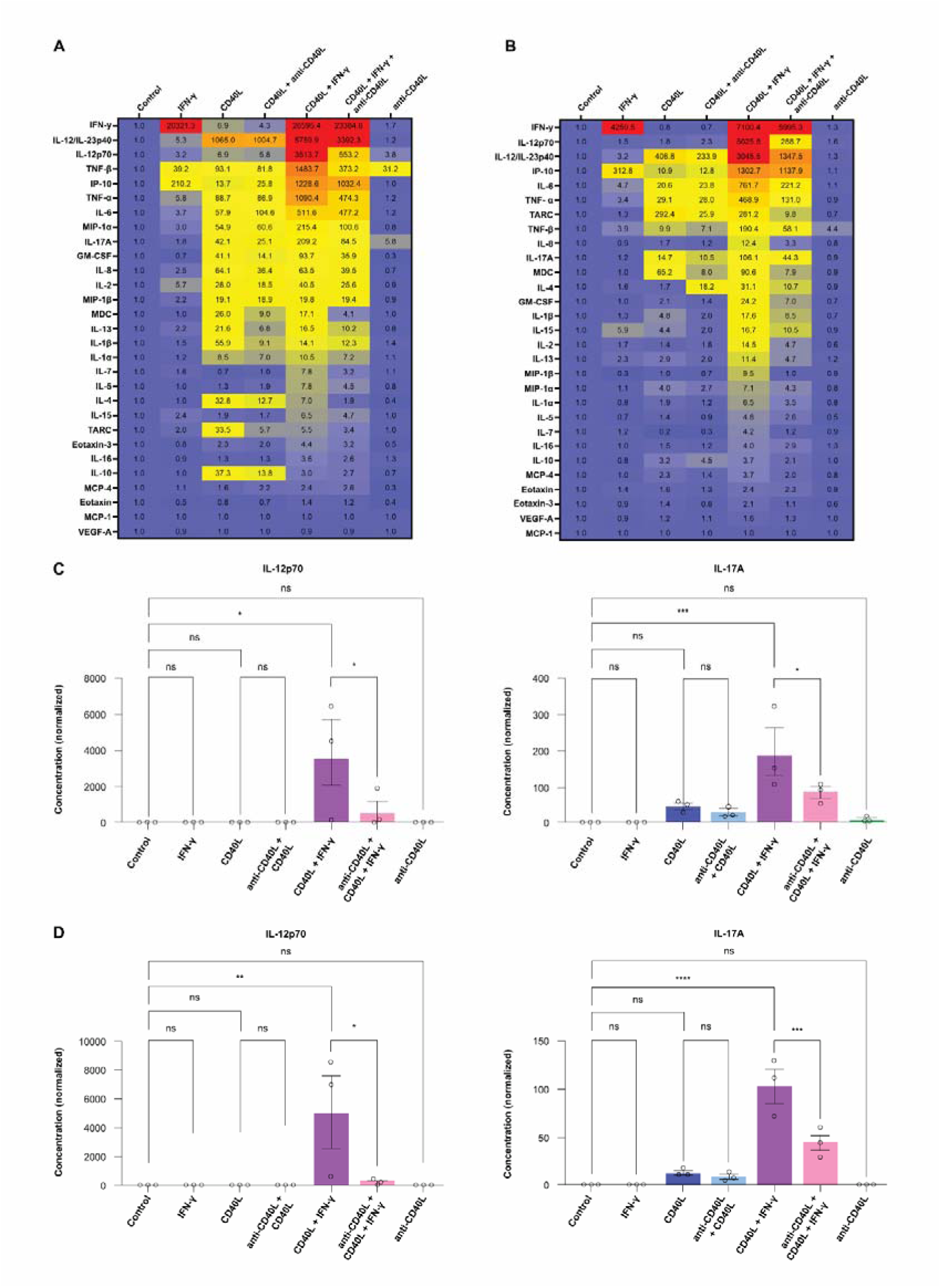
Robust inflammatory activation by co-stimulation with CD40L and IFN-γ is attenuated with anti-CD40L in tri-cultures. A Heat map -24 h after co-stimulation with CD40L and IFN-γ. B Heat map -5 days after co-stimulation with CD40L and IFN-γ. C IL-12p70 and IL-17A -24 h after co-stimulation with CD40L and IFN-γ. D IL-12p70 and IL-17A 5 days after co-stimulation with CD40L and IFN-γ. Cytokine concentrations in the medium over hiPSC-derived tri-cultures co-stimulated with CD40L, IFN-γ, and/or an anti-CD40L tool antibody (10 µg/mL). Numbers displayed in the heatmap in A denote average cytokine concentrations across experiments, normalized to the vehicle control within experiment. Bar charts in C and D also display average cytokine concentrations across experiments, normalized to vehicle control within experiment, with each dot representing an individual experiment (n=3 independent experiments). Statistical significance within the heatmap and bar chart are indicated by * *p*-value≤0.05, ** *p*-value≤0.01, *** *p*-value≤0.001, and **** *p*-value≤0.0001. Data are presented as mean±SEM. Source data are provided as a Source Data file. CD40L, cluster of differentiation 40 ligand; hiPSC, human induced pluripotent stem cell; IFN-γ, interferon-gamma; IL, interleukin; SEM, standard error of the mean; TNF-α, tumor necrosis factor-alpha.

Interestingly, IL-12p70 release was selectively elicited by co-stimulation with CD40L and IFN-γ, but not by CD40L or IFN-alone (Fig. 4C and 4D). The resulting cytokine signatures closely resembled those observed in microglia mono-cultures (Fig. 1E and Fig. S4). To evaluate astrocytic responses, we co-stimulated astrocyte mono-cultures with CD40L and/or IFN-γ and measured the cytokine release at 24 h. Astrocyte mono-cultures showed minimal responses (Fig. S9A and S9B), further suggesting that microglia are the main drivers of CD40L-mediated inflammation and neuronal damage.

### CD40L-enriched gene signature prevalent in PPMS perilesional tissue

CD40L-mediated microglial activation has been implicated in MS and autoimmune CNS inflammation^14,15^, but the role of the pathway in progressive disease is less well understood. Furthermore, the specific microglial gene programs and molecular pathways that can be reversed by anti-CD40L therapeutic intervention remain poorly defined. To address this critical knowledge gap, we performed single-nucleus RNA sequencing on perilesional brain tissue from five patients with PPMS (male, n=2; female, n=3; mean age, 57 ± 9.6 years; post-mortem interval [PMI], 14.2 ± 7.9 h) and on normal-appearing white matter (NAWM) from six age-matched and sex-matched healthy controls (male, n=3; female, n=3; mean age, 68.8 ± 11.5 years; PMI: 16.7 ± 4.4 h) (Fig. 5A). We identified all major CNS cell types, as well as B cells and T cells, in both groups (Fig. 5B-D). As expected, T cell and B cell numbers were higher in PPMS tissue than in NAWM (Fig. 5C and Fig. S10). Notably, we observed high CD40 expression in the microglia cluster (Fig. 5E), supporting the relevance of CD40-mediated signaling in PPMS perilesional microglia.

**Fig. 5.**
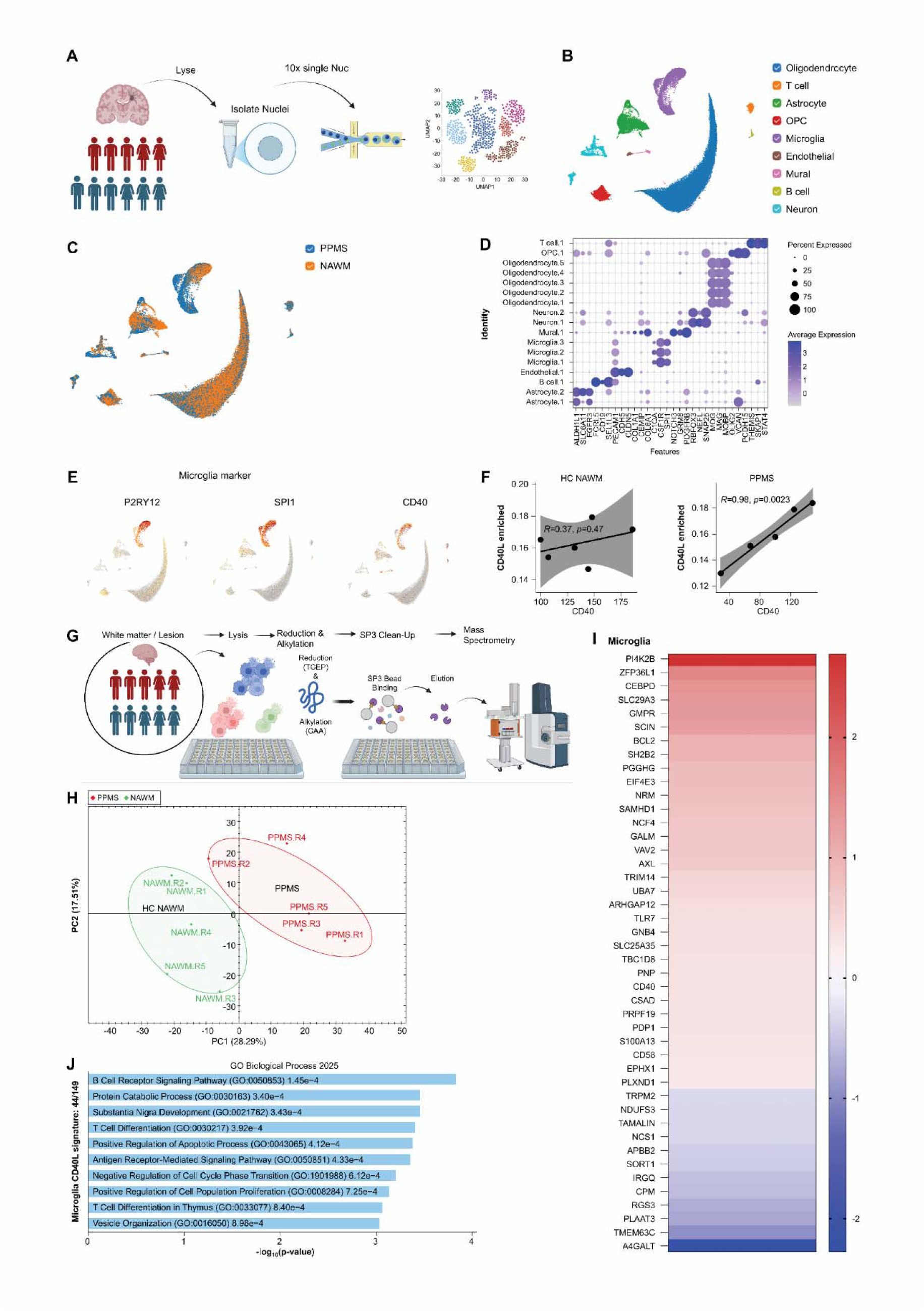
CD40L signature present in PPMS perilesional tissue. A Overview of number and sex of patients and control and schema for single-nucleus RNA sequencing (PPMS n=5 and NAWM n=6). B UMAP separated by cell type. C UMAP separated by disease condition: NAWM or PPMS. D Identification of cell type and cluster by expression of cell-type-specific markers. E UMAP expression of microglia markers *P2RY12* and *SPI1* and *CD40*, showing targeted upregulation of CD40 in the microglia cluster. F Correlation of enrichment for CD40L signature genes to CD40 expression in NAWM (n=6) and PPMS (n=5) patients (Pearson correlation). G Overview of number and sex of patients and control and schema for DIA-MS (PPMS n=5, NAWM n=5). H PCA plot of NAWM and PPMS samples. I Heatmap of significantly expressed proteins that overlapped with microglia CD40L enriched gene signature samples. J Number of genes from the microglia CD40L signature that were significantly dysregulated at the protein level in PPMS tissue and pathway analysis. CD40, cluster of differentiation 40; CD40L, cluster of differentiation 40 ligand; DIA-MS, direct data-independent acquisition mass spectrometry; NAWM, normal-appearing white matter; *P2RY12*, purinergic receptor *P2Y12*; PCA, principal component analysis; PPMS, primary progressive multiple sclerosis; RNA, ribonucleic acid; *SPI1*, Spi-1 proto-oncogene (also known as PU.1); UMAP, Uniform Manifold Approximation and Projection.

We defined a gene signature from the hiPSC-microglia bulk RNA sequencing dataset by identifying any genes that were significantly dysregulated (adjusted *p*-value*<*0.05, |fold change|<0.8 or >1.2) when exposed to CD40L and IFN-γ and then fully recovered to vehicle control levels with anti-CD40L treatment (Fig. 2D). We then performed single-nucleus gene set enrichment analysis on microglia from the NAWM controls and PPMS perilesional tissue samples, calculating enrichment scores for the CD40L-reversible gene expression and correlating these with CD40 expression levels using Pearson correlation. We observed a significant correlation (R=0.98, *p*-value=0.0023) in the PPMS group but no correlation in the NAWM group (R=0.37, *p*-value =0.47) (Fig. 5F), suggesting that CD40L activation may be present in progressive MS perilesional microglia. For further investigation, we performed proteomic profiling on brain tissue from an overlapping cohort.

We performed single-shot direct data-independent acquisition mass spectrometry (DIA-MS) on flash-frozen white matter lesions from five patients with PPMS (male, n=3; female, n=2; mean age: 57.2 ± 9.3 years; PMI: 12.3 ± 7.1 h) and five NAWM samples from age-matched neurologically healthy controls (male, n=2; female, n=3; mean age: 68.6 ± 12.9 years; PMI: 17.7 ± 4.2 h) (Fig. 5G). PCA revealed a clear separation between PPMS lesions and NAWM controls (PC1: 28.3% variance; PC2: 17.5% variance; Fig. 5H). Notably, of the 149 genes in the microglial CD40L-responsive signature, 44 showed significant dysregulation (adjusted *p*-value<0.05, |fold change|<0.8 or >1.2) at the protein level (Fig. 5I). Pathway analysis of these dysregulated proteins again highlighted immune and inflammatory responses, including B cell activation and T cell differentiation (Fig. 5J), suggesting that CD40L inhibition within the CNS may limit disease progression by suppressing microglial and B cell activation.

## DISCUSSION

MS is characterized by progressive neurodegeneration driven by compartmentalized CNS inflammation that persists despite effective peripheral immune modulation^24^. While CD40L-expressing T cells activate CD40-expressing B cells and may contribute to MS pathology^4,25^, the CD40/CD40L axis may contribute to the smoldering neuroinflammation by activating microglia in progressive disease. Our tri-culture does not recapitulate all the complexity of CD40L signaling in the brain and, we do not know whether in MS patients CD40L-CD40 signaling between T cells and microglia is cell-cell contact mediated or whether soluble CD40L (sCD40L) is cleaved from the T cell surface and acts at a distance from its cellular source. sCD40L has also been implicated in MS pathology. In the CSF of patients with RRMS, sCD40L levels have been positively correlated with the number of gadolinium-positive lesions^26^, suggesting that both membrane-bound and soluble forms of CD40L might contribute to disease activity, albeit through distinct routes. The relative contribution of these two modes of signaling may shift across disease phases, reflecting underlying differences in T cell-microglia spatial dynamics^24^. In early MS, T cells and microglia are in close proximity within perivascular spaces and the CNS parenchyma, which may enable direct contact-dependent CD40/CD40L signaling that drives TNF-α release, chemokine production, and neuronal injury. In progressive MS, the BBB is less permeable and T cells become increasingly compartmentalized within meningeal, perivascular, and lesion-border spaces, with reduced parenchymal infiltration^24,27^. Under these conditions, microglia at the lesion border may be activated in part through soluble mediators, including shed sCD40L derived from border-zone T cells, which could amplify proinflammatory responses at a distance. This represents one of several contributors to progressive pathology alongside other CNS-intrinsic processes, such as sustained intrathecal cytokine production, which has itself been shown to drive MS-like cortical demyelination and neurodegeneration independent of peripheral immune infiltration^28^. Notably, plasma sCD40L levels are elevated in SPMS relative to non-progressive benign MS and RRMS^29^, and elevated sCD40L has been detected in the CSF of patients with MS, although its levels across specific MS subtypes remain to be fully established^30^. Additionally, tissue-resident memory CD8 T cells that persist in the parenchyma show markers of local reactivation, potentially through antigen re-encounter via microglial antigen presentation^27^. Regardless of the source of CD40L, CD40 signaling in microglia represents a robust inflammatory node that might contribute to neurodegeneration in progressive MS lesions and warrants consideration as a therapeutic target in the disease.

While our findings do not delineate the two sources of CD40L, hiPSC-derived microglia mono-cultures demonstrate that CD40L activation induced cell clustering, inflammatory cytokine release, and eventually cell death. These effects were markedly exacerbated by co-administration of IFN-γ, a proinflammatory cytokine produced by activated T cells. IFN-γ alone did not induce microglial clustering or cell death, and its effects on cytokine release were modest compared to those elicited by CD40L or CD40L and IFN-γ co-administration. Previous studies in mice have shown that IFN-γ can prime microglia by increasing CD40 expression^14,18^, while our findings show that CD40L was required to induce a full inflammatory response. Among the cytokines analyzed, IL-12, TNF-α, and IL-6 exhibited the greatest increases following stimulation with CD40L or co-stimulation with CD40L and IFN-γ. Changes in these cytokines following stimulation *in vitro* partially align with cytokine changes found in MS. IL-12p40 is elevated in CSF in RRMS and correlates with disease activity^31^ and TNF-α levels are increased in serum, CSF, and active brain lesions of patients with MS, with peak levels observed during clinical exacerbations^32,33^. IL-6 is elevated in the CSF of patients with MS and is associated with clinical and radiological disease activity, including the presence of contrast-enhancing MRI lesions^34-37^. These cytokine changes in MS could be caused by a number of different upstream mechanisms but could partially converge on CD40-CD40L signaling in the periphery and CNS.

We found that CD40 activation in hiPSC microglia drove a striking clustering behavior *in vitro*. Microglial clustering has previously been observed in early and late stage MS brain tissue^38-40^ and in animal models of MS near leaky blood vessels, but whether these observed changes are CD40 mediated has not been examined^41^. Upon CD40L engagement of CD40 microglia may aggregate to form a local proinflammatory niche, analogous to the immune cell clustering observed during antigen presentation in peripheral lymphoid tissues. This interpretation is supported by the concurrent upregulation of antigen presentation-related transcripts, including *LAMP3*, *HLA-F*, *IDO1*, and *CCR7*, in CD40L-stimulated microglia as well as pathway enrichment for B cell activation and proliferation. Temporally, clustering preceded the onset of cell death, suggesting it represents an upstream activation event rather than a consequence of dying cells. In the current study, stimulation of microglia with either CD40L alone or co-stimulation with CD40L and IFN-γ elicited unique morphological responses. Within 24 h, CD40L alone caused microglia to adopt a rod-like morphology, similar to that reported in MS subpial gray matter lesions^42^. In contrast, co-stimulation with CD40L and IFN-γ induced a hyper-ramified morphology, within the same timeframe. These distinct morphologies further highlight the divergence of microglial responsiveness to CD40L alone versus CD40L with IFN-γ. Bulk RNA sequencing revealed that co-stimulation with CD40L and IFN-γ presents a greater-than-additive effect. Transcriptomic pathway enrichment analyses identified multiple pathways related to B cell activation and proliferation. In progressive MS, B cells invade both the brain border tissues and the CNS parenchyma and lesions^24,43,44^, where CD40L-activated microglia may release factors that further increase B cell proliferation and activation. In addition to B cell activation, we observed significant dysregulation of lysosomal and phagocytic function pathways. Recent transcriptomic and spatial profiling studies have demonstrated substantial dysregulation of phagocytic, lysosomal, lipid-metabolic, and cholesterol-handling pathways in lesion-associated microglia and macrophages, resulting in lipid-laden inflammatory phenotypes that are linked to chronic active lesion expansion in progressive MS^22,45^. Whether CD40 mediated phagocytic activation of microglia is helpful or harmful for microglia to drive clearance and metabolism of different extracellular substrates is unknown but should be further investigated.

Employing hiPSC-derived tri-culture system, we demonstrated that CD40L and IFN-γ combined elicit responses in microglia that are capable of mediating neuronal injury and neurite degeneration in human cells. In this system, co-stimulation with CD40L and IFN-γ induced progressive neuronal dysfunction, increased cell death, and reduced neurite density. This neuronal dysfunction appears to be microglia dependent and was not observed with CD40L alone and only minimally observed with IFN-γ alone; consistently, neither stimulus alone reduced neurite density. These results imply that after activation of the microglia by CD40L and IFN-γ, microglia produce factors that impact neuron health and/or physically interact with neurons to drive neuronal death. The effect could also be explained by an indirect effect on astrocytes, which are activated by microglia-derived cytokines to cause neurotoxicity^46^. We found that the CD40L/IFN-γ driven neuronal death was partially blocked by a necroptosis inhibitor. This suggests that the cell death could be driven by TNF-α, interferons, or other cytokines which activate the necroptotic pathway^47^, but more detailed studies would be needed to figure out how this signaling works. Anti-CD40L treatment only partially prevents neuronal death, which was surprising given how well that it inhibited cytokine release in the microglia mono-cultures. It is possible that a residual amount of CD40 signaling was sufficient to still drive neuron death, and that higher concentrations of the inhibitor could block this effect.

Several limitations of this study warrant consideration. First, while hiPSC-derived models recapitulate key aspects of human CNS biology, they do not fully replicate the complexity of the *in vivo* CNS microenvironment, including BBB integrity, chronic demyelination, and the full spectrum of peripheral immune cell infiltration. Although our tri-culture system incorporates neurons, astrocytes, and microglia, it lacks oligodendrocytes and endothelial cells, both of which may influence microglial activation state and contribute to MS pathology. Second, the human tissue analyses focused on perilesional tissue from patients with PPMS, but the involvement of CD40L pathway activation in chronic inactive lesions or other MS subtypes, including RRMS or SPMS, remains to be established. Nevertheless, the CD40L+IFN-γ-induced transcriptional signature identified in hiPSC-derived microglia is enriched in PPMS perilesional tissue, correlates strongly with microglial CD40 expression, and is recapitulated at the protein level in PPMS lesions, validating its strong translational relevance. Comparative analyses across disease stages and subtypes may reveal whether CD40L-targeted therapies have broad applicability or are most beneficial in specific patient populations.

## MATERIALS AND METHODS

### Human *in vitro* microglia culture

#### Cell culture procedures

hiPSC-derived microglia (BrainXell; BX-0900-32) or GFP-expressing microglia (BrainXell; BX-0901-30) were seeded at 4–5×10^4^ cells/well in poly-D-lysine (PDL)-coated 96-well plates and maintained in Dulbecco’s Modified Eagle Medium (DMEM)/Nutrient Mixture F-12 (DMEM/F-12) (Gibco, 11330-032) supplemented with N2 supplement (1×; Gibco, 17502048), GlutaMAX (0.5×; Gibco, 35050-061), B27 supplement (0.5×; Gibco, 17504044), minimum essential medium non-essential amino acids (MEMNEAA) (1×; Gibco, 11140-050), chemically defined lipid concentrate (0.5%; Gibco, 11905-031), ascorbic acid 2-phosphate (65 μg/mL; Sigma, A8960), 2-mercaptoethanol (0.00084%), antibiotic-antimycotic (1×; Gibco, 15240062), transforming growth factor-beta 1 (TGF-β1) (2 ng/mL), macrophage colony-stimulating factor (M-CSF) (20 ng/mL), and IL-34 (100 ng/mL). Half-medium changes were performed every 2–3 days, and cells were cultured for ≥4 days prior to analyses.

#### Live-cell imaging

IFN-γ was included as a co-stimulus as it has been reported to modify microglial responsiveness to CD40L via upregulation of CD40 expression and potentially other synergistic mechanisms^19^. Hoechst stained or GFP expressing microglia were imaged on an Opera Phenix™ High Content Screening System (Revvity, HH14001000, USA) with a 20× water-immersion objective. Cells were imaged 24 h after treatment with either recombinant human IFN-γ (100 U/mL; R&D Systems, 285-IF-100/CF), recombinant human CD40L (5 μg/mL; R&D Systems, 6420-CLB-050), and/or an anti-CD40L tool antibody (10 μg/mL). Nearest-neighbor nuclear distance was quantified for Hoechst-stained microglia using Harmony software (version 5.1; Revvity). Nuclear visualization and quantification of microglial clustering was performed using Hoechst staining.

#### Cell death assay

Cell death was quantified by DRAQ7-positive nuclei using an Incucyte® SX5 Live-Cell Analysis System (Sartorius, 4816, Germany). Five days after plating, cultures were treated with DRAQ7 reagent and the following: either recombinant human IFN-γ (100 U/mL; R&D Systems, 285-IF-100/CF), recombinant human CD40L (5 μg/mL) (R&D Systems, 6420-CLB-050), an anti-CD40L antibody (Sanofi, internal compound) (1, 10, or 100 μg/mL), RIPK1 inhibitor GSK547 (10 μM), RIPK3 inhibitor GSK872 (3 μM), ferroptosis inhibitor liproxstatin-1 (100 nM), and/or the NLRP3 inflammasome inhibitor MCC950 (5 μM). Imaging began 1 h after treatment and continued for up to 7 days.

#### Cytokine/chemokine immunoassays

To assess cytokine and chemokine secretion, hiPSC-derived microglia were cultured as described above (*Section: Cell culture procedures*). Five days after plating, cells were treated with recombinant human IFN-γ (100 U/mL), recombinant human CD40L (5 μg/mL), and/or an anti-CD40L antibody (1 or 10 μg/mL). After 24 h, cell culture supernatants were collected and assayed using the MESO QuickPlex SQ 120MM (Meso Scale Diagnostics, AI1AA-0, USA) multiplex immunoassays, including the Human Proinflammatory Panel 1 (K15049D), Human Chemokine Panel 1 (MSD, K15047D), and Human Cytokine Panel 1 (MSD, K15050D). Supernatants were collected at 24 h post stimulation to avoid confounding effects of cell death, which begins at approximately 36 h under co-stimulation conditions.

#### Bulk RNA sequencing and analysis

hiPSC-derived microglia were seeded at 2.5×10^5^ cells/well in PDL-coated 24-well plates and maintained as described above (*Section: Cell culture procedures*). Five days after plating, cells were treated with either recombinant human IFN-γ (100 U/mL), recombinant human CD40L (5 μg/mL), and/or an anti-CD40L antibody (10 μg/mL). After 12 h, cells were lysed in RLT buffer (350 μL per well; QIAGEN, 79216), subjected to a brief freeze−thaw cycle to enhance lysis, and stored at −80 °C for RNA analysis.

RNA library preparation and sequencing were performed by Fulgent Genetics. Total RNA was processed using the Illumina TruSeq Stranded mRNA Library Prep Kit according to the manufacturer’s instructions. Briefly, polyadenylated RNA was isolated using oligo-dT magnetic beads, fragmented, and converted to first-strand complementary DNA (cDNA) using random hexamer primers and reverse transcriptase. Second-strand cDNA synthesis incorporated dUTP to preserve strand specificity. After end repair, A tailing, and adapter ligation, libraries were PCR amplified, quality checked TapeStation 4200 (Agilent Technologies, G2991BA, Germany), quantified (Qubit), and then sequenced on a NovaSeq 6000 (Illumina, 20012850, USA) (150 bp paired end reads).

Raw reads were imported into QIAGEN OmicSoft Studio (version 12.8) and aligned to GRCh38 using the OSA4 aligner with default parameters (≤2 mismatches; multi-mappers excluded). Counts were assigned to OmicSoft GenCode v47 models and normalized as RPKM. Differential expression analysis was performed using a General Linear Model in OmicSoft Studio with Benjamini–Hochberg false discovery rate (FDR) correction. Genes with |log2 fold-change |>1 and FDR<0.05 were considered significant.

### Human *in vitro* astrocyte culture

#### Cell culture procedures

hiPSC-derived astrocytes (Fujifilm, C1037) were seeded at a density of 5×10^4^ cells/well in PDL-coated 96-well plates and cultured in DMEM/F-12 medium supplemented with heat-inactivated fetal bovine serum (FBS) (2%) and N2 supplement (Gibco, 17502-048). Experiments were performed 5 days after cell plating.

#### Immunoassays

To assess cytokine and chemokine secretion, hiPSC-derived astrocytes were prepared, and MSD multiplex immunoassays were performed as described above for microglia culture. To evaluate astrocytic contributions, astrocyte mono-cultures were co-stimulated with CD40L (5 μg/mL) and/or IFN-γ (100 U/mL) and cytokine release was measured at 24 h.

### Human *in vitro* tri-culture

#### Cell culture procedures

hiPSC-derived tri-cultures were generated in a stepwise manner. Astrocytes (Fujifilm, C1037) were first seeded on Day 0 at a density of 1.5×10^4^ cells/well onto Matrigel-coated 96-well plates (Corning, 354277) and maintained in DMEM/F-12 with heat-inactivated FBS (2%) and N2 supplement. On Day 1, motor neurons (Fujifilm, C1048) were seeded onto the astrocyte layer at 3.5×10^4^ cells/well and maintained in iCell Neural Base Medium (Fujifilm, M1010) with Neural

Supplement A (Fujifilm, M1032) and Nervous System Supplement (Fujifilm, M1031). On Day 14, microglia (BrainXell, BX-0900-32) or GFP-expressing microglia (BrainXell, BX-0901-30) were seeded at 2.0×10^4^ cells/well into wells containing both astrocytes and motor neurons. After seeding, cells were maintained in tri-culture medium consisting of iCell Neuron Medium with Microglia Supplement A (Fujifilm, M1036), TGF-β1 (2 ng/mL), M-CSF (20 ng/mL), and IL-34 (100 ng/mL). Cells consistently underwent a half-medium change every 2–3 days, and tri-cultures were maintained for at least 4 days following the addition of microglia before being used for analyses.

All hiPSC-derived cells from Fujifilm are considered under the global MSA agreement established between Sanofi and FCDI and have been reviewed and approved by the Sanofi Ethics Department. The parental iPSC lines used to generate these products (identified as 01434 and 01279) are registered in the Human Pluripotent Stem Cell Registry (hPSCReg), where independent third-party ethical review and certification of the informed consent process (ICF) were completed, confirming that appropriate donor consent had been obtained. The accrediting authorities and corresponding approval numbers are Rush University IACUC (13-034 and 14-072) for line 01279 and NICHD-NIH (N01-HD-4-2865) for line 01434.

#### Neuronal calcium imaging

Incucyte Neuroburst Lentivirus (Sartorius, 4736; synapsin promoter-driven orange-fluorescent calcium indicator) was added on culture Day 7 at 10 µL/well. On Day 8, culture wells were washed thrice, followed by a full-medium change; thereafter, culture maintenance proceeded as described above. Calcium imaging was performed daily, starting on the day of treatment, using the Incucyte^®^ SX5 Live-Cell Analysis System. Quantification of neuronal activity was conducted using the Incucyte^®^ Neuronal Activity Analysis Software Module, reporting active neuron count (i.e., active object count), calcium transient intensity (i.e., mean intensity), and calcium transient synchrony (i.e., mean correlation). All Neuroburst fluorescence intensity traces were normalized as Δ*F*/*F*_0_ prior to metric calculation. Cultures were treated with the indicated stimuli, following the procedures and concentrations described in the preceding section.

#### Live-cell imaging

To track temporal microglial morphological changes, GFP-expressing microglia were imaged longitudinally on Days 1, 3, 5, 7, and 10 post stimulation. Active neuronal populations were quantified in parallel via calcium imaging to correlate microglial dynamics with neuronal loss. Live-cell imaging of GFP-expressing hiPSC-derived microglia within tri-cultures was performed using an Opera Phenix™ High Content Screening System with a 20× water-immersion objective. Imaging was conducted on Days 1, 3, 5, 7, and 10 after treatment with either recombinant human IFN-γ (100 U/mL) and/or recombinant human CD40L (5 μg/mL).

#### Immunocytochemistry

Tri-cultures were fixed with paraformaldehyde (4%) 14 days after treatment with either recombinant human IFN-γ (100 U/mL), recombinant human CD40L (5 μg/mL), and/or an anti-CD40L antibody (10 or 100 μg/mL). Fixed cultures were then incubated for 1 h at room temperature in antibody-dilution buffer (5% bovine serum albumin and 0.1% Triton-X 100 in PBS) and subsequently incubated overnight at 4 °C with the following primary antibodies: beta-III tubulin (1:100; Millipore Sigma, MAB1637), IBA1 (1:500; Abcam, ab5076), and GFAP (1:500; Millipore Sigma, AB5541). The following day, cultures were washed with 0.1% Tween-20/PBS and then incubated for 1 h at room temperature with the following secondary antibodies: anti-mouse AlexaFluor 647 (Thermo Fisher Scientific, A31571), anti-goat AlexaFluor 555 (Abcam, A21432), and anti-chicken AlexaFluor 488 (Abcam, A78948). After additional washes with 0.1% Tween-20/PBS followed by PBS, cultures were imaged using an Opera Phenix™ High Content Screening System with a 20× water-immersion objective. Image staining areas were quantified using Harmony software (version 4.9).

#### Cell death assay

Cell death in tri-cultures was assessed by quantifying DRAQ7-stained nuclei using an Incucyte^®^ SX5 Live-Cell Analysis System; treatments followed the mono-culture conditions.

#### Immunoassays

To assess cytokine and chemokine secretion, hiPSC-derived tri-cultures were cultured as described above. Supernatants were collected at 24 h and 5 days post stimulation to capture the immediate cytokine response and the inflammatory milieu coinciding with the onset of active neuron loss, respectively. Supernatants collected 24 h or 5 days after treatment were assayed via MSD panels (K15049D, K15047D, K15050D).

### Single nuclei isolation, sequencing, and analysis

#### Single-nucleus sample preparation

Brain tissue from PPMS patients and healthy control were obtained from the NIH NeuroBioBank. Frozen brain tissue (50 mg) samples were processed using the Chromium Nuclei Isolation Kit with RNase inhibitor (10× Genomics, 1000494). After the final wash, nuclei were resuspended in 300 µL wash buffer, stained with 4,6-diamidino-2-phenylindole (DAPI) (Thermo Fisher Scientific, EN62248), and counted. Nuclei were further purified via magnetic-activated cell sorting (MACS) with anti-nucleus microbeads (Miltenyi Biotec, 130-132-997). Purified nuclei were extracted into 300 µL of separation buffer, stained again with DAPI, and re-counted.

#### Single-nucleus RNA sequencing

Nuclei were processed with the Chromium GEM-X Single Cell 3’ v4 kit (10× Genomics; 1000691, 1000690, 1000215) and processed following the manufacturer’s protocols. All samples (n=11) were processed in parallel: cDNA was amplified for 11 cycles during initial amplification, and sample-index PCR was performed for 14 cycles. Libraries were sequenced on an Illumina NovaSeq 6000 (S4, 200 cycle kit; Illumina, 20028313) and processed with Cell Ranger 8 against the GRCh38 2020 A reference.

### Proteomics by Liquid chromatography-tandem mass spectrometry (LC–MS/MS)

Proteomic analysis was performed using single-pot, solid-phase-enhanced sample processing (SP3)^48^. Briefly, brain samples were lysed in 2% sodium dodecyl sulfate (SDS), 300 mM triethylammonium bicarbonate (TEAB), 300 mM NaCl, 8 M urea with protease/phosphatase inhibitors, followed by probe sonication at 12% amplitude. Lysates were clarified by centrifugation (21,130 × *g*, 10 min, 4 °C), and protein concentration was quantified using the Pierce BCA assay. Proteins were reduced and alkylated using tris(2-carboxyethyl)phosphine (TCEP; 20 mM) and chloroacetamide (CAA; 40 mM) at 60 °C for 30 min, respectively. Sera Mag SpeedBeads (1:1 hydrophilic:hydrophobic; Cytiva) were added (5 μL), ethanol was adjusted to 50% (v/v), and samples were incubated 30 min at room temperature (600 rpm). Beads were washed with 80% ethanol and acetonitrile (ACN), then resuspended in 50 μL of 50 mM TEAB with trypsin (1:50, w/w). After overnight digestion (37 °C), peptides were eluted with LC–MS grade water and 200 mM TEAB, pooled, dried (vacufuge), acidified with 1% formic acid, and loaded onto Evotips.

All mass spectrometry data were acquired using a timsTOF HT coupled to an EvosepOne LC system via a CaptiveSpray nano-electrospray source. Peptides were separated on a C18 reversed-phase column (15 cm × 75 μm ID, 1.6 μm; IonOpticks) containing an integrated CaptiveSpray emitter. Runs were performed using the 20 SPD Whisper protocol. Data independent acquisition parallel accumulation serial fragmentation (diaPASEF) was performed using twenty-one 25-Da precursor double-window scans spanning 475−1000 m/z and 1/k_0_ of 0.85−1.27 V.s/cm^2^. Library-free direct DIA+ analysis was performed using Spectronaut^®^ 19.8 (Biognosys). Raw data were searched against the SwissProt human database (taxon ID: 9606; 20,626 entries, January 2022). Database search criteria followed standard DIA+ settings: tryptic digestion with up to two missed cleavages, carbamidomethylation of cysteine as a fixed modification, oxidation of methionine as a variable modification, and a precursor Q-value (FDR) cut-off of 0.01.

### Statistical analysis

Data are presented as mean±standard error of the mean (SEM). *N* represents the number of technical replicates within individual experiments and the number of independent experiments as specified, in the figure legends. Statistical analyses were performed using GraphPad Prism software (version 10). Statistical significance is indicated * *p*-value≤0.05, ** *p*-value≤0.01, *** *p*-value≤0.001, and **** *p*-value≤0.0001 and was determined by one-way analysis of variance (ANOVA) with Sidak’s post hoc test.

## Contributors

Gregory S. Wirak, Sean Ryan, and Timothy Hammond conceived and designed the studies. Dimitry Ofengeim and Timothy Hammond contributed to all aspects of the work, including conceptualization, data curation, formal analysis, investigation/methodology, project administration, supervision, validation, visualization, and writing, review and editing. Gregory S. Wirak and Sean Ryan contributed to conceptualization, data curation, project administration, resources, software, supervision, validation, visualization, and writing, review and editing. Anna Blazier contributed to formal analysis, supervision, validation, visualization, and writing, review and editing. Ivan Nyarko-Danquah contributed to conceptualization, data curation, formal analysis, supervision, validation, and visualization. Dongyu Liu contributed to formal analysis and supervision, validation, and visualization. Bo Yang contributed to conceptualization, data curation, formal analysis, and writing, review and editing. Roopali Gandhi contributed to conceptualization, data curation, investigation/methodology, and writing, review and editing. Biljana Djukic contributed to conceptualization, data curation, and writing, review and editing. Jose Vela contributed to conceptualization, data curation, and investigation/methodology. Shameer Khader and Hendrik Wesseling contributed to project administration, resources, software, supervision, validation, and visualization. Michael LaMorte contributed to conceptualization and data curation. Bailin Zhang and Franck Rapaport contributed to project administration, resources, and software. All authors have read and approved the final manuscript.

|  | Conceptualization, data curation | Formal analysis | Investigation/methodology | Project administration, resources, and software | Supervision, validation, and visualization | Writing - review and editing |
|---|---|---|---|---|---|---|
| **Gregory S. Wirak** | Y | Y | Y | Y | Y | Y |
| **Sean Ryan** | Y | Y | Y | Y | Y | Y |
| **Anna Blazier** |  | Y |  |  | Y |  |
| **Dongyu Liu** |  | Y |  |  | Y |  |
| **Ivan Nyarko-Danquah** | Y | Y |  |  | Y |  |
| **Michael LaMorte** | Y |  |  |  |  |  |
| **Shameer Khader** |  |  |  | Y | Y |  |
| **Jose Vela** | Y |  | Y |  |  |  |
| **Bo Yang** | Y | Y |  |  | Y | Y |
| **Hendrik Wesseling** |  |  |  | Y | Y |  |
| **Bailin Zhang** |  |  |  | Y |  |  |
| **Franck Rapaport** |  |  |  | Y |  |  |
| **Roopali Gandhi** | Y |  | Y |  | Y | Y |
| **Biljana Djukic** | Y |  |  |  |  | Y |
| **Dimitry Ofengeim** | Y |  |  |  |  |  |
| **Timothy Hammond** | Y | Y | Y | Y | Y | Y |

## Data sharing

All data used to support the findings of this study are included within the article, and raw data are available from the corresponding author. Further details on data-sharing criteria, eligible studies, and process for requesting access are available <u>online</u>.

## Acknowledgments

Sanofi funded the study. Anti-CD40L is being developed by Sanofi. Tissue specimens were obtained from the NIH Brain and Tissue Repository-California, Human Brain and Spinal Fluid Resource Center, VA West Los Angeles Medical Center, Los Angeles, California, which is supported in part by National Institutes of Health and the US Department of Veterans Affairs. The authors thank Samuel Oliver, PharmD from Sanofi for his contribution to manuscript planning, review, and coordination and Deepa Warrier, PhD, and Shailaja Mahajan, PhD, from Sanofi, for providing medical writing support.

