## Supplementary Figures S1 to S10 for "CD40 ligand and IFN-γ induce proinflammatory activity and microglia-mediated neurodegeneration in human stem cell models"

### **Fig S1: Morphology of GFP-expressing hiPSC-derived microglia**

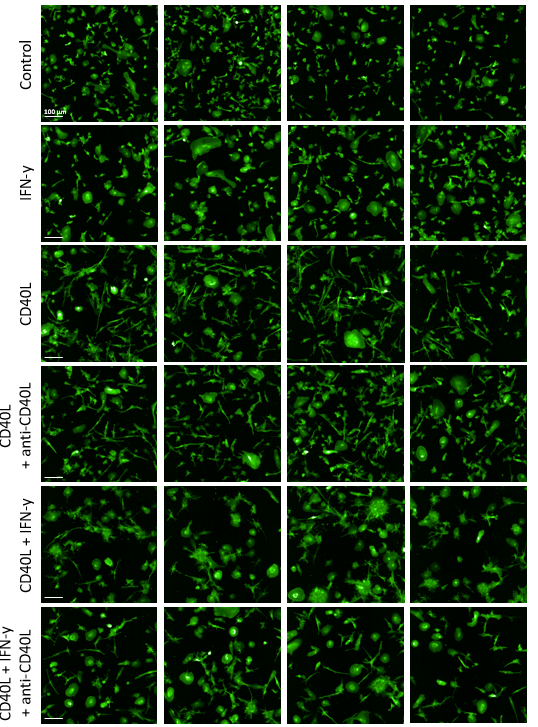

Confocal fluorescence imaging of live GFP-expressing hiPSC-derived microglia, 24 h after stimulation with CD40L, IFN-γ, and/or an anti-CD40L antibody.

CD40L, cluster of differentiation 40 ligand; GFP, green fluorescent protein; IFN-γ, interferon-gamma; hiPSC, human induced pluripotent stem cell.

### **Fig S2: Morphology and distribution of hiPSC-derived microglia**

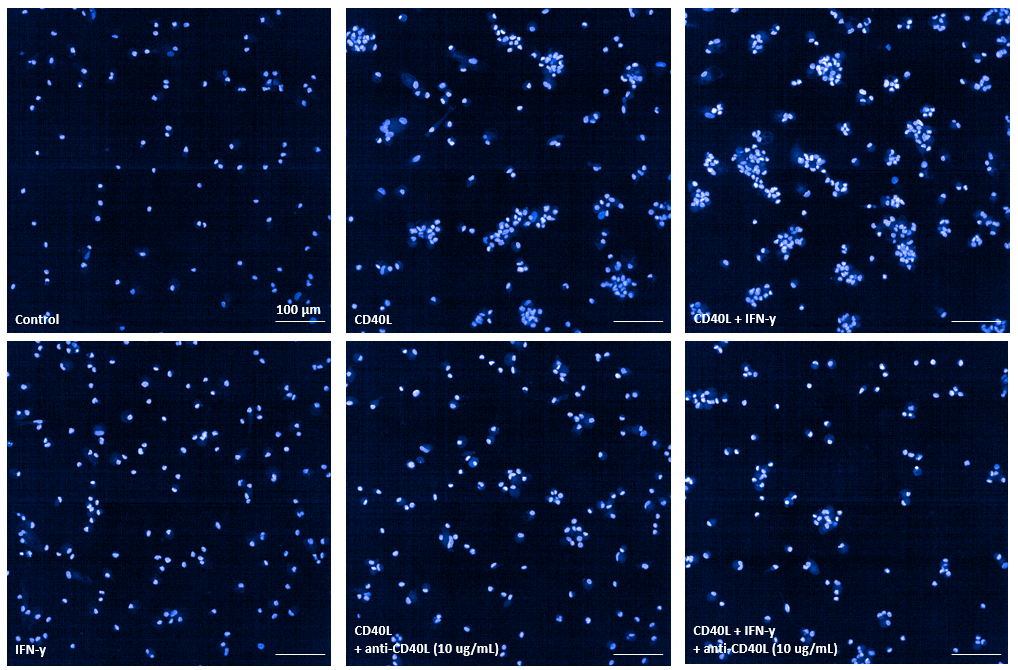

Confocal fluorescence imaging of Hoechst-stained nuclei in live microglia, 24 h after stimulation with CD40L, IFN-γ, and/or an anti-CD40L antibody.

CD40L, cluster of differentiation 40 ligand; hiPSC, human induced pluripotent stem cell; IFN-γ, interferon-gamma.

### **Fig S3: Microglial death with various cell death inhibitors**

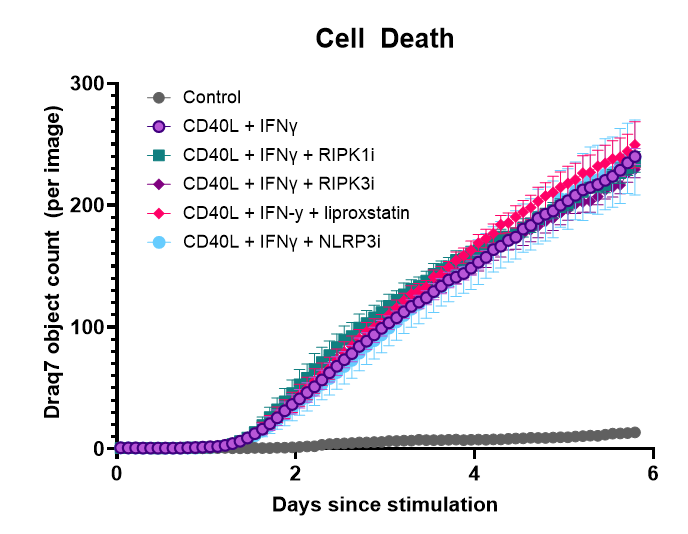

Time course of dead cell counts in DRAQ7^TM^-stained hiPSC-derived microglia cultures, seven days after stimulation with CD40L, IFN-γ, and/or various inhibitors of cell death mediators (i.e. inhibitors RIPK1, RIPK3, and NLRP3 and the ferroptosis inhibitor liproxstatin) (n=3 independent experiments). Data are presented as mean values ±SEM. Source data are provided as a Source Data file.

CD40L, cluster of differentiation 40 ligand; DRAQ7, deep red anthraquinone 7; hiPSC, human induced pluripotent stem cell; IFN-γ, interferon-gamma; NLRP3, NOD-like receptor protein 3; RIPK1, receptor-interacting protein kinase 1; RIPK3, receptor-interacting protein kinase 3; SEM, standard error of the mean.

### **Fig S4: Cytokine release profile of hiPSC-derived microglia**

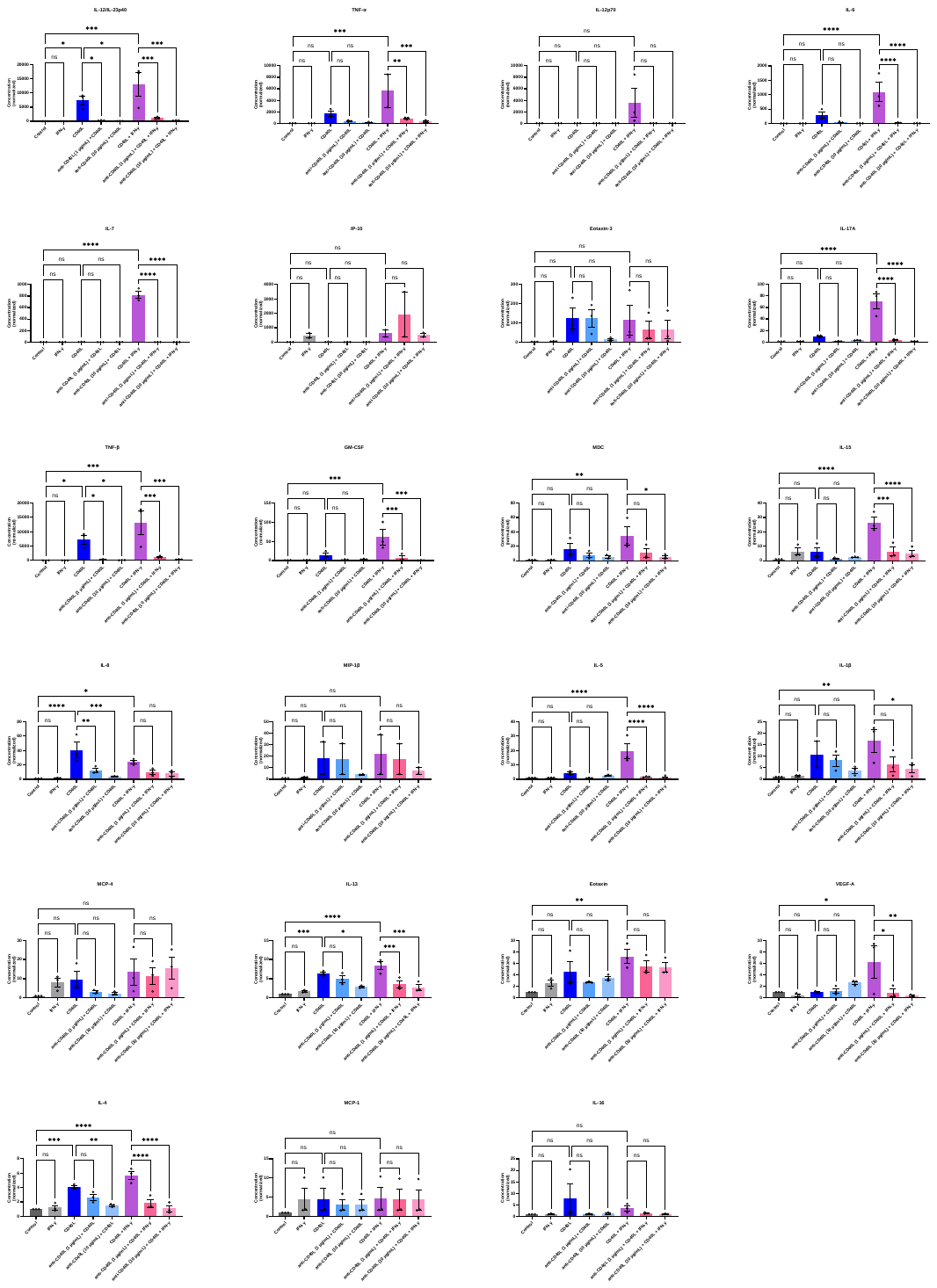

Cytokine concentrations in the medium over microglia cultures stimulated with CD40L, IFN-γ, and/or an anti-CD40L antibody. Bar charts display average cytokine concentrations across experiments, normalized to vehicle control within experiments, with each dot representing an individual experiment (n=3 separate experiments). **(A)** is the same heatmap as in **Fig. 1E**, expanded to include indicators of statistical significance. Statistical significance within the heatmap and bar chart are indicated by * *p*-value≤0.05, ** *p*-value≤0.01, *** *p*-value≤0.001, and **** *p*-value≤0.0001. Data are presented as mean values ±SEM. Source data are provided as a Source Data file.

CD40L, cluster of differentiation 40 ligand; GM‑CSF, granulocyte‑macrophage colony‑stimulating factor; hiPSC, human induced pluripotent stem cell; IL, interleukin; IFN-γ, interferon-gamma; IP‑10, interferon‑gamma–induced protein 10 (CXCL10); MCP, monocyte chemoattractant protein; MDC, macrophage‑derived chemokine; MIP, macrophage inflammatory protein; SEM, standard error of the mean; TNF‑α, tumor necrosis factor alpha; VEGF‑A, vascular endothelial growth factor‑A.

### **Fig S5: Metrics of neuronal activity in hiPSC-derived tri-cultures 48 h after stimulation with CD40L and IFN-γ**

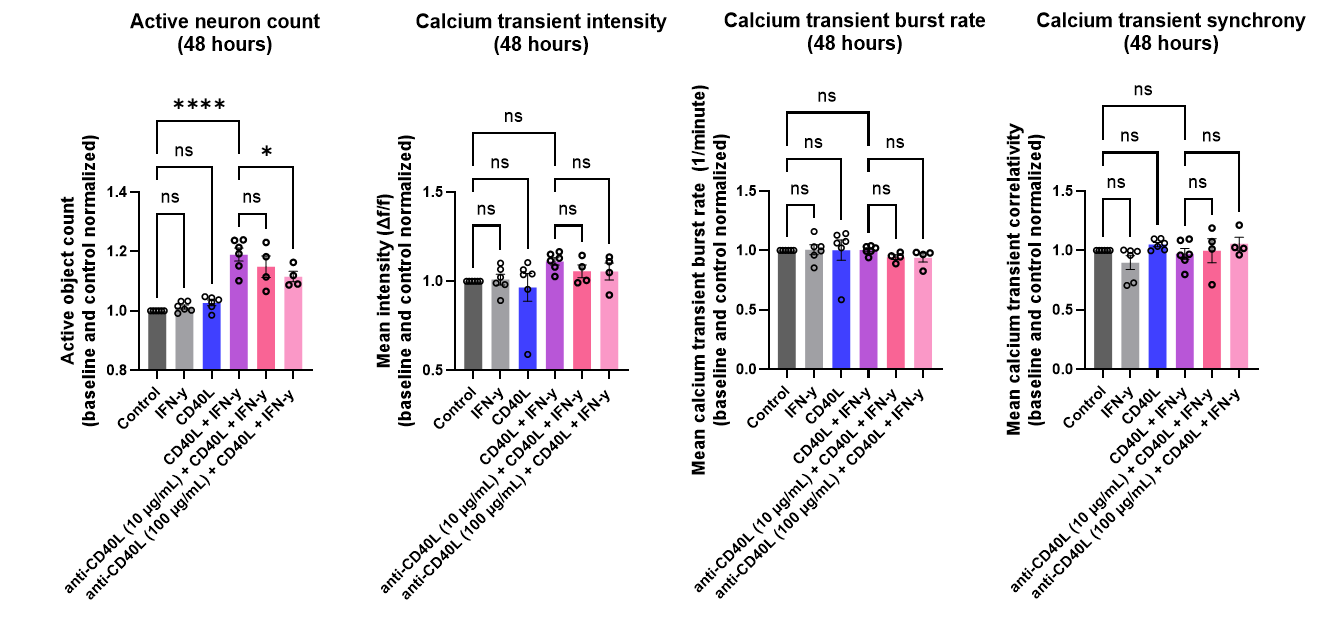

Active neuron counts, calcium transient intensities, calcium transient burst rates, and calcium transient correlativity (graphs, left to right) within hiPSC-derived tri-cultures of microglia, astrocytes, and neurons stimulated with CD40L, IFN-γ, and/or an anti-CD40L tool antibody (n=4-6 individual experiments). Neuronal activity was captured via neuron-specific calcium imaging 48 h after stimulation. Values shown are normalized to within-experiment controls. Statistical significance is indicated by * *p*-value≤0.05, ** *p*-value≤0.01, *** *p*-value≤0.001, and **** *p*-value≤0.0001. Data are presented as mean values ± SEM. Source data are provided as a Source Data file.

CD40L, cluster of differentiation 40 ligand; hiPSC, human induced pluripotent stem cell; IFN-γ, interferon-gamma; ns, not significant; SEM, standard error of the mean.

### **Fig S6: Cellular morphologies and distributions within hiPSC-derived tri-cultures conditions**

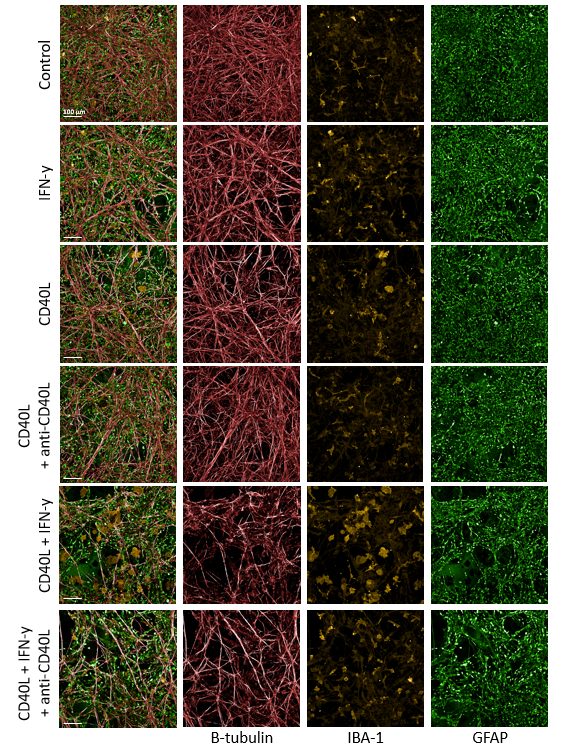

Fluorescent immunostaining of markers of neurons (beta-III tubulin), microglia (IBA1), and astrocytes (GFAP) in tri-cultures stimulated with CD40L, IFN-γ, and/or an anti-CD40L antibody, 14 days after stimulation.

CD40L, cluster of differentiation 40 ligand; GFAP, glial fibrillary acidic protein; hiPSC, human induced pluripotent stem cell; IBA1, ionized calcium‑binding adapter molecule 1; IFN‑γ, interferon‑gamma.

**Fig S7: Cellular morphologies and distributions of hiPSC-derived microglia**

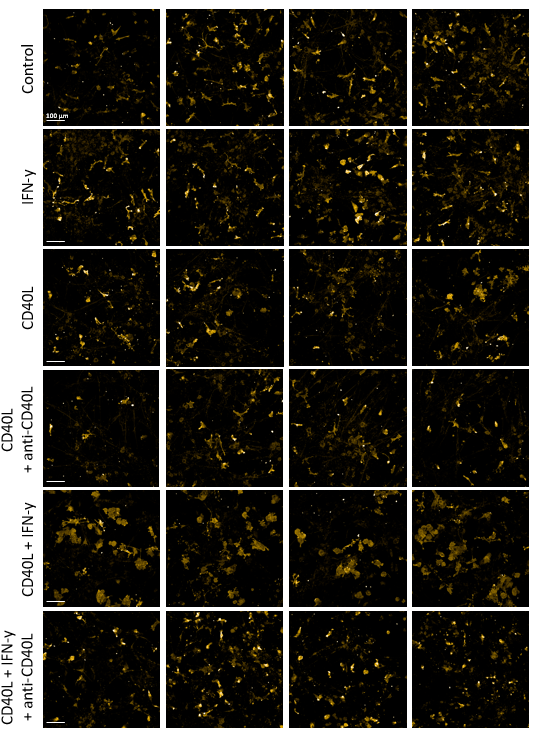

Fluorescent immunostaining of a marker of microglia (IBA1) in tri-cultures stimulated with CD40L, IFN-γ, and/or an anti-CD40L antibody, 14 days after stimulation.

CD40L, cluster of differentiation 40 ligand; hiPSC, human induced pluripotent stem cell; IBA1, ionized calcium‑binding adapter molecule 1; IFN‑γ, interferon‑gamma.

### **Fig S8: Shifting morphology of GFP-expressing hiPSC-derived microglia, in parallel with progressive neuronal insult**

**A**

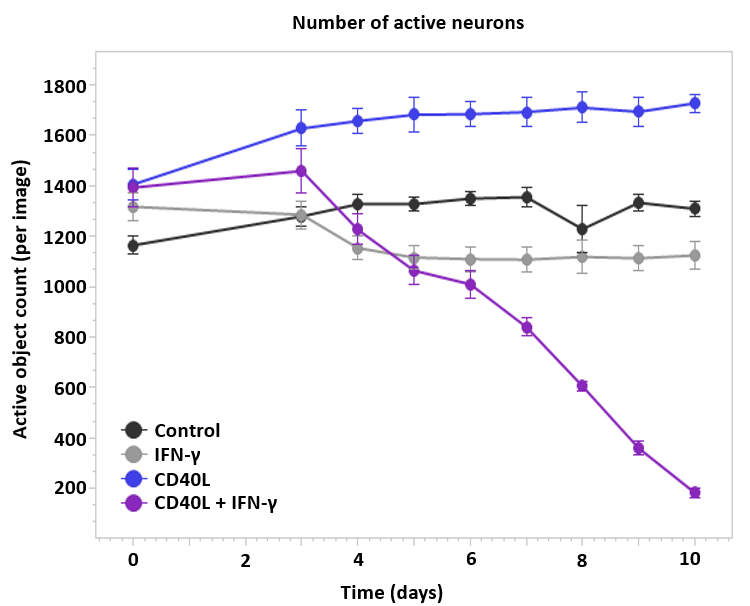

**B**

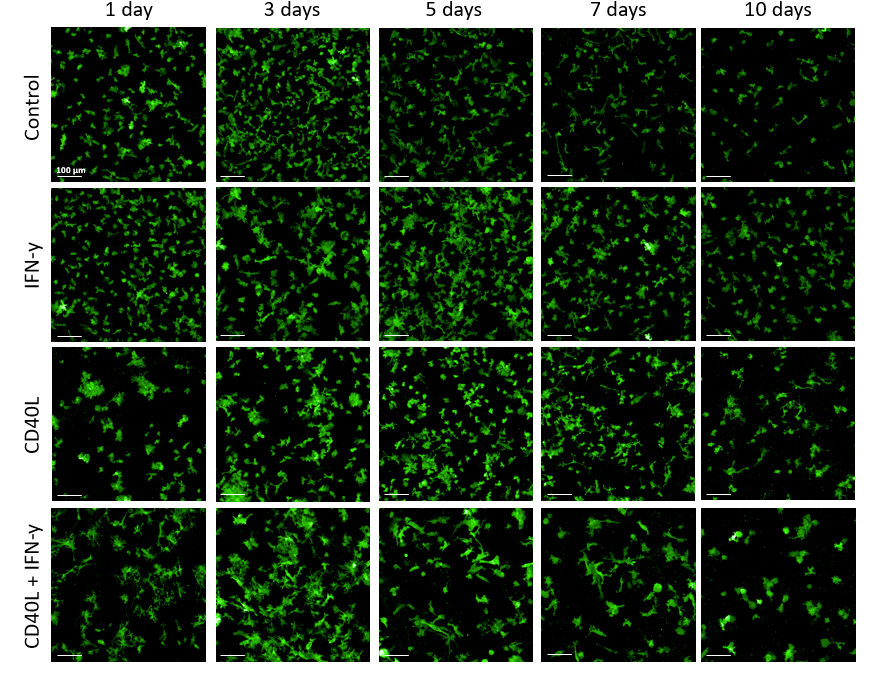

**A** Time course of active neuron counts within hiPSC-derived tri-cultures of microglia, astrocytes, and neurons, up to 10 days after stimulation with CD40L and/or IFN-γ (n=6 technical replicates). Displayed are the average raw counts per image with one timelapse image acquired per culture plate well. Neuronal activity was captured via neuron-specific calcium imaging every 24 h. **B** Confocal fluorescence imaging time course of the live GFP-expressing hiPSC-derived microglia within the tri-cultures used to generate data for **A**. Data are presented as mean values ± SEM. Source data are provided as a Source Data file.

CD40L, cluster of differentiation 40 ligand; GFP, green fluorescent protein; hiPSC, human induced pluripotent stem cell; IFN‑γ, interferon‑gamma; SEM, standard error of the mean.

**B**

### **Fig S9: Cytokine release profile of hiPSC-derived astrocytes**

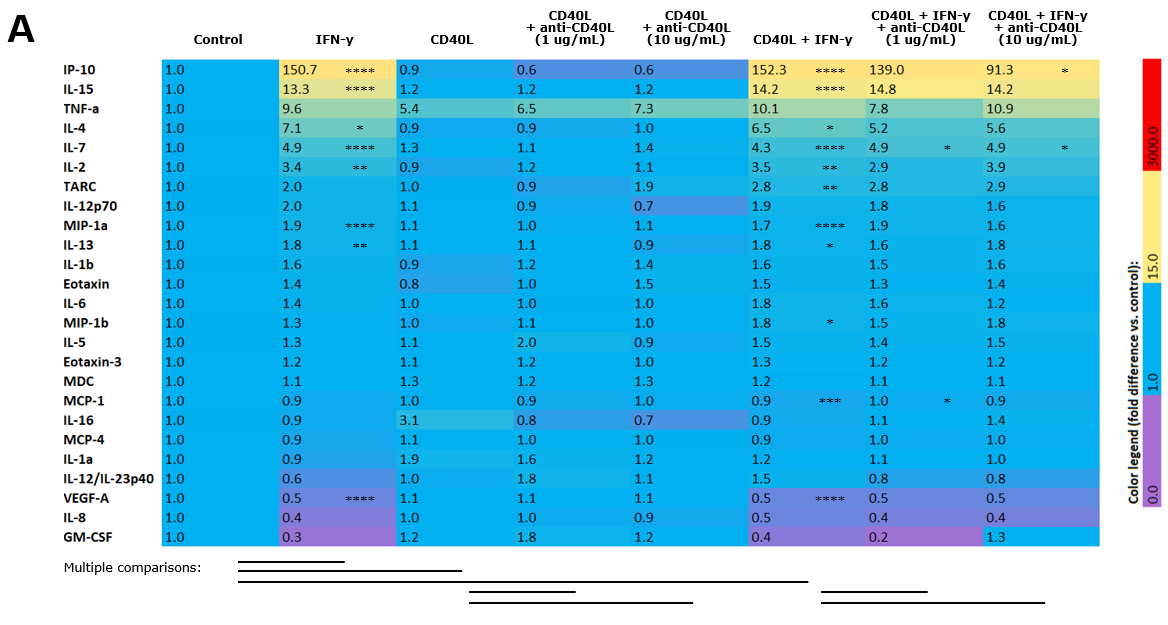

### **Fig S9: Cytokine release profile of hiPSC-derived astrocytes**

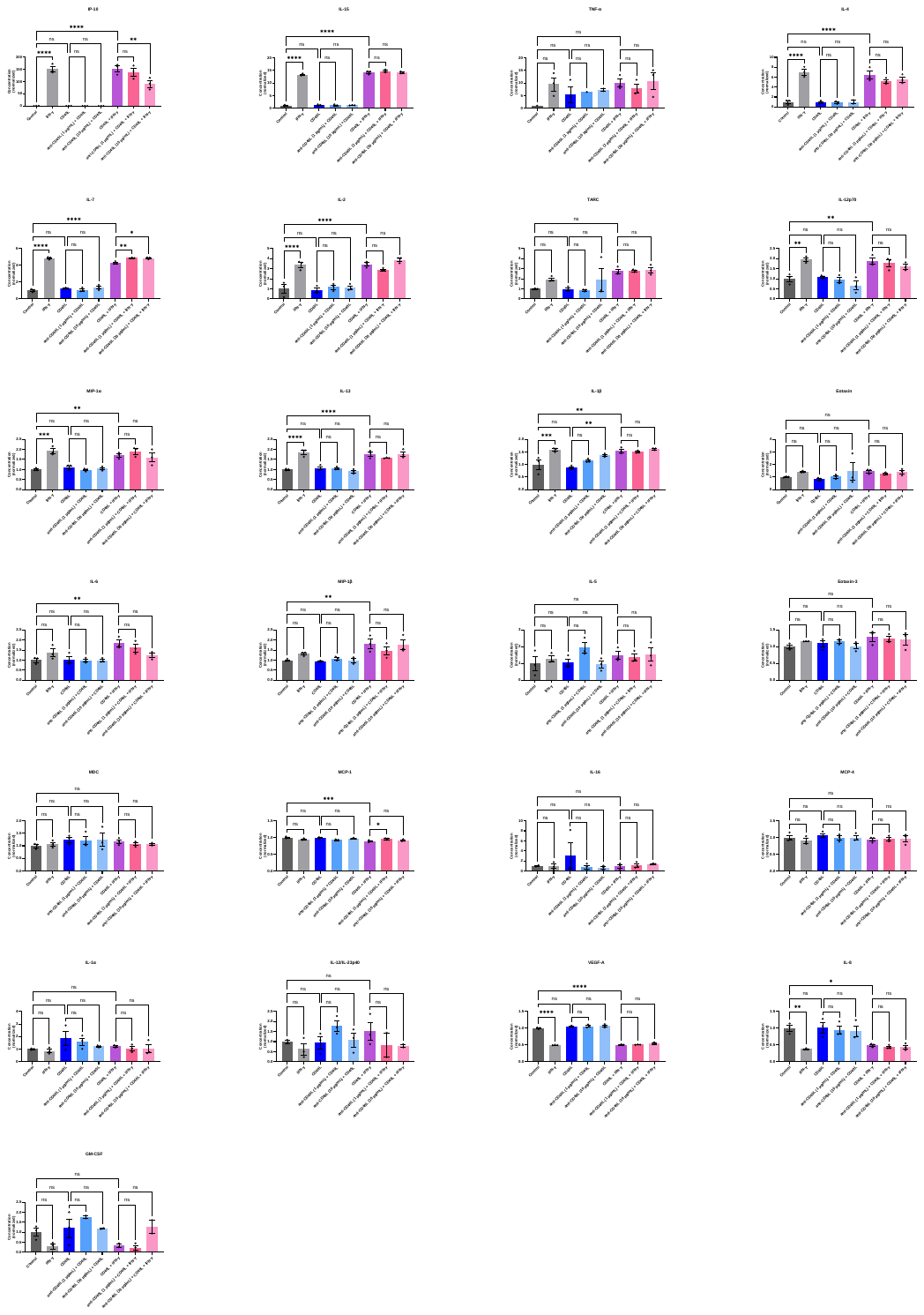

**B**

Cytokine concentrations in the medium over astrocyte cultures stimulated with CD40L, IFN-γ, and/or an anti-CD40L tool antibody. Numbers displayed on the heatmap in **A** denote average cytokine concentrations across replicates, normalized to vehicle control. Bar charts in **B** also display average cytokine concentrations across replicates, normalized to vehicle control, with each dot representing an individual replicate (n = 3 technical replicates). Statistical significance within the heatmap and bar chart are indicated by * *p*-value≤0.05, ** *p*-value≤0.01, *** *p*-value≤0.001, and **** *p*-value≤0.0001. Data are presented as mean values ± SEM. Source data are provided as a Source Data file.

CD40L, cluster of differentiation 40 ligand; GM‑CSF, granulocyte‑macrophage colony‑stimulating factor; hiPSC, human induced pluripotent stem cell; IFN-γ, interferon-gamma; IL, interleukin; IP‑10, interferon‑gamma–induced protein 10 (CXCL10); MCP, monocyte chemoattractant protein; MDC, macrophage‑derived chemokine; MIP, macrophage inflammatory protein; TARC, thymus and activation‑regulated chemokine; SEM, standard error of the mean; TNF‑a, tumor necrosis factor alpha; VEGF‑A, vascular endothelial growth factor‑a.

### **Fig S10: PPMS Single Nucleus RNA sequencing cell counts**

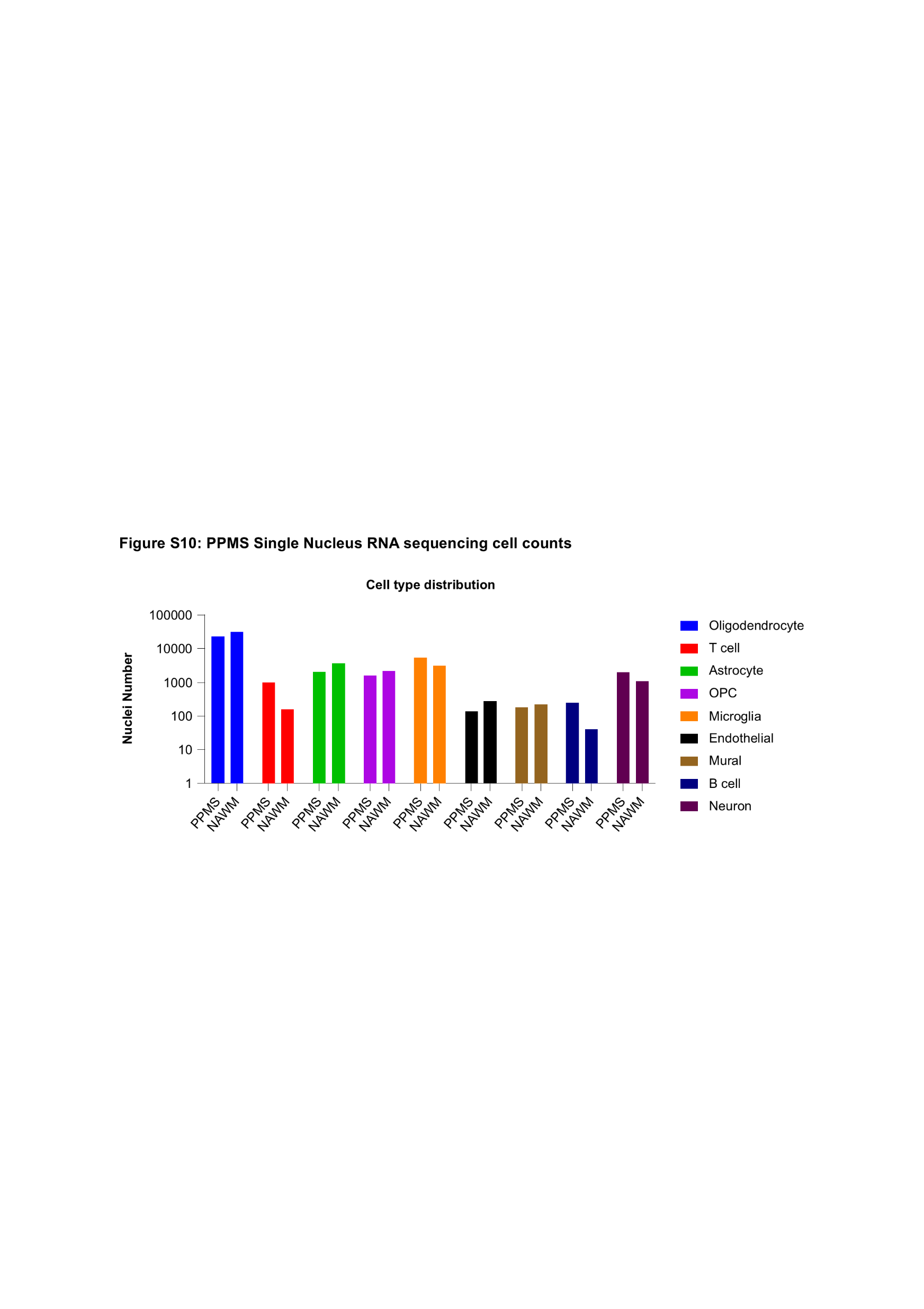

Number of nuclei per cell type identified by single-nucleus RNA sequencing in perilesional brain tissue from patients with PPMS (n=5) and NAWM from healthy controls (n=6). Bars represent absolute nucleus counts per cell type for each condition.

NAWM, normal-appearing white matter; OPC, oligodendrocyte precursor cells; PPMS, primary progressive multiple sclerosis; RNA, ribonucleic acid.
